# A novel ion conducting route besides the central pore in an inherited mutant of G-protein-gated inwardly rectifying K^+^ channel

**DOI:** 10.1101/2021.08.18.456735

**Authors:** I-Shan Chen, Jodene Eldstrom, David Fedida, Yoshihiro Kubo

## Abstract

G-protein-gated inwardly rectifying K^+^ (GIRK; Kir3.x) channels play important physiological roles in various organs. Some of the disease-associated mutations of GIRK channels are known to induce loss of K^+^ selectivity but their structural changes remain unclear. In this study, we investigated the mechanisms underlying the abnormal ion selectivity of inherited GIRK mutants. By the two-electrode voltage-clamp analysis of GIRK mutants heterologously expressed in *Xenopus* oocytes, we observed that Kir3.2 G156S permeates Li^+^ better than Rb^+^, while T154del or L173R of Kir3.2 and T158A of Kir3.4 permeate Rb^+^ better than Li^+^, suggesting a unique conformational change in the G156S mutant. Applications of blockers of the selectivity filter (SF) pathway, Ba^2+^ or Tertiapin-Q (TPN-Q), remarkably increased the Li^+^-selectivity of Kir3.2 G156S but did not alter those of the other mutants. In single-channel recordings of Kir3.2 G156S expressed in mouse fibroblasts, two types of events were observed, one attributable to a TPN-Q sensitive K^+^ current and the second a TPN-Q resistant Li^+^ current. The results show that a novel Li^+^ permeable and blocker-resistant pathway exists in G156S in addition to the SF pathway. Mutations in the pore helix (PH), S148F and T151A, also induced high Li^+^ permeation. Our results demonstrate that the mechanism underlying the loss of K^+^ selectivity of Kir3.2 G156S involves formation of a novel ion permeation pathway besides the SF pathway, which allows permeation of various species of cations.

## Introduction

G-protein-gated inwardly rectifying K^+^ (GIRK) channels are stimulated by the activation of G-protein-coupled receptors (GPCRs) to regulate heartbeat, neuronal excitability and hormone secretion (Hibino et al., 2010). There are four types of GIRK subunits (Kir3.1, Kir3.2, Kir3.3, and Kir3.4) which form homotetramers of Kir3.2 or heterotetramers of multiple combinations in different tissues (Hibino et al., 2010; Krapivinsky et al., 1995; Kubo, Reuveny, Slesinger, Jan, & Jan, 1993; Lesage et al., 1994), and gene mutations encoded by GIRK channels have been found to cause diseases in some patients (Zangerl-Plessl, Qile, Bloothooft, Stary-Weinzinger, & van der Heyden, 2019). For example, deletion of Thr152, or mutations of Gly154 to Ser or Leu171 to Arg in the human Kir3.2 channel (encoded by *KCNJ6*), cause Keppen-Lubinsky syndrome (KPLBS) or KPLBS-like disorder (Horvath et al., 2018; Masotti et al., 2015), and G151R, T158A or L168R mutations in the human Kir3.4 channel (encoded by *KCNJ5)* cause aldosterone-producing adenomas (APA) (Boulkroun et al., 2012; Choi et al., 2011). These amino acid residues are conserved among GIRK subtypes. Some of the inherited mutations of these residues have been reported to cause loss of K^+^ selectivity, resulting in disorder of cell function and cell death (Choi et al., 2011; Horvath et al., 2018; Navarro et al., 1996; Scholl et al., 2012; Slesinger et al., 1996).

K^+^ selectivity is achieved by a highly conserved amino acid sequence TV(I)GYG which forms the selectivity filter (SF) in K^+^ channels (Zagotta, 2006). Mutations in the SF sequence of K^+^ channels alter ion selectivity (Heginbotham, Lu, Abramson, & MacKinnon, 1994). An ion pair, conserved in the Kir channel family, a glutamic acid located in the pore helix (PH) and an arginine located in the extracellular loop behind the SF, is known to contribute to stabilization of the pore structure and maintaining ion selectivity (Yang, Yu, Jan, & Jan, 1997). Mutations of other residues in the PH, such as Ser148 or Glu152 of mouse Kir3.2, also impair the K^+^ selectivity (Chen et al., 2019; Yi, Lin, Jan, & Jan, 2001). Mutations located in transmembrane domains (TMs) outside the SF also alter ion selectivity and residues located at the central cavity can also influence the ion selectivity through interaction with the permeating ions (Bichet, Grabe, Jan, & Jan, 2006; Bichet et al., 2004; Matamoros & Nichols, 2021; Yi et al., 2001). Structures of various types of K^+^ channel have been identified by Cryo-EM and X-ray crystallography (Long, Campbell, & Mackinnon, 2005; Miller & Long, 2012; W. Wang & MacKinnon, 2017; Whorton & MacKinnon, 2011). However, the structural changes induced by mutations at different regions of the channel which alter the ion selectivity still remain to be elucidated.

Previous studies using homology modeling have predicted structural changes to GIRK channels induced by disease associated mutations as follows: (A) mutations in the SF (human Kir3.2 T152del or Kir3.2 G154S) impair the stabilization of K^+^ interaction with the backbone of the SF (Masotti et al., 2015); (B) the L171R mutation in the TM2 of human Kir3.2 limits the SF movement by the formation of a hydrogen bond with Glu148 in the PH (Horvath et al., 2018), while the L168R mutation at the corresponding position of human Kir3.4 is relevant to the interaction with the side chain of Tyr of the GYG motif in the SF (Choi et al., 2011); (C) the T158A mutation in the extracellular loop of human Kir3.4 weakens the stability of the structure by eliminating the hydrogen bonds between the extracellular loop, PH, and TM1 (Choi et al., 2011). However, no further experimental data have been provided to support these hypotheses.

In the present study, we investigated the mechanisms underlying the abnormal ion selectivity of these inherited mutations in Kir3.2 and Kir3.4 channels by introducing mutations at the corresponding positions in mouse Kir3.2 or rat Kir3.4. Using electrophysiological recordings in *Xenopus* oocytes, we demonstrate that most of these mutations induce a conformational change which allows increased permeation of Rb^+^ or Cs^+^, while the G156S mutation of the Kir3.2 channel (corresponding to human Kir3.2 G154S) induces a different conformational change which allows better permeation of Li^+^ or Na^+^ over Rb^+^ or Cs^+^. Applications of pore blockers of GIRK channels increase the Li^+^-selectivity of Kir3.2 G156S but do not influence that of the other mutants. Single-channel recordings of the G156S mutant in mouse fibroblasts show that two types of ion conducting events, which respectively reflect K^+^ current via the SF pathway and Li^+^ current via a novel ion permeation pathway, exist in the same recording. Mutations of amino acid residues located in the PH behind the SF also gave rise to the Li^+^-permeable pathway outside of the SF route. Our results reveal a novel mechanism underlying the loss of K^+^-selectivity of Kir3.2 G156S that involves formation of a novel ion permeation pathway in addition to the conventional SF route.

## Results

### Different disease associated mutations of GIRK channels show diverse ion selectivity

To characterize the ion selectivity changes of GIRK channels induced by inherited mutations, we constructed these mutants using mouse Kir3.2 or rat Kir3.4 channels (Figure 1a). These mutants were heterologously expressed in *Xenopus* oocytes and the ionic currents were recorded by two-electrode voltage-clamp. We first recorded the K^+^ current with an extracellular solution containing 96 mM K^+^, and then replaced the K^+^ with other cations. The order of their ionic size is Li^+^ < Na^+^ < K^+^ < Rb^+^ < Cs^+^ < methylammonium^+^ (MA^+^) < tetramethylammonium^+^ (TMA^+^) < tetraethylammonium^+^ (TEA^+^) < n-methyl-d-glucamine^+^ (NMDG^+^).

**Figure 1.**
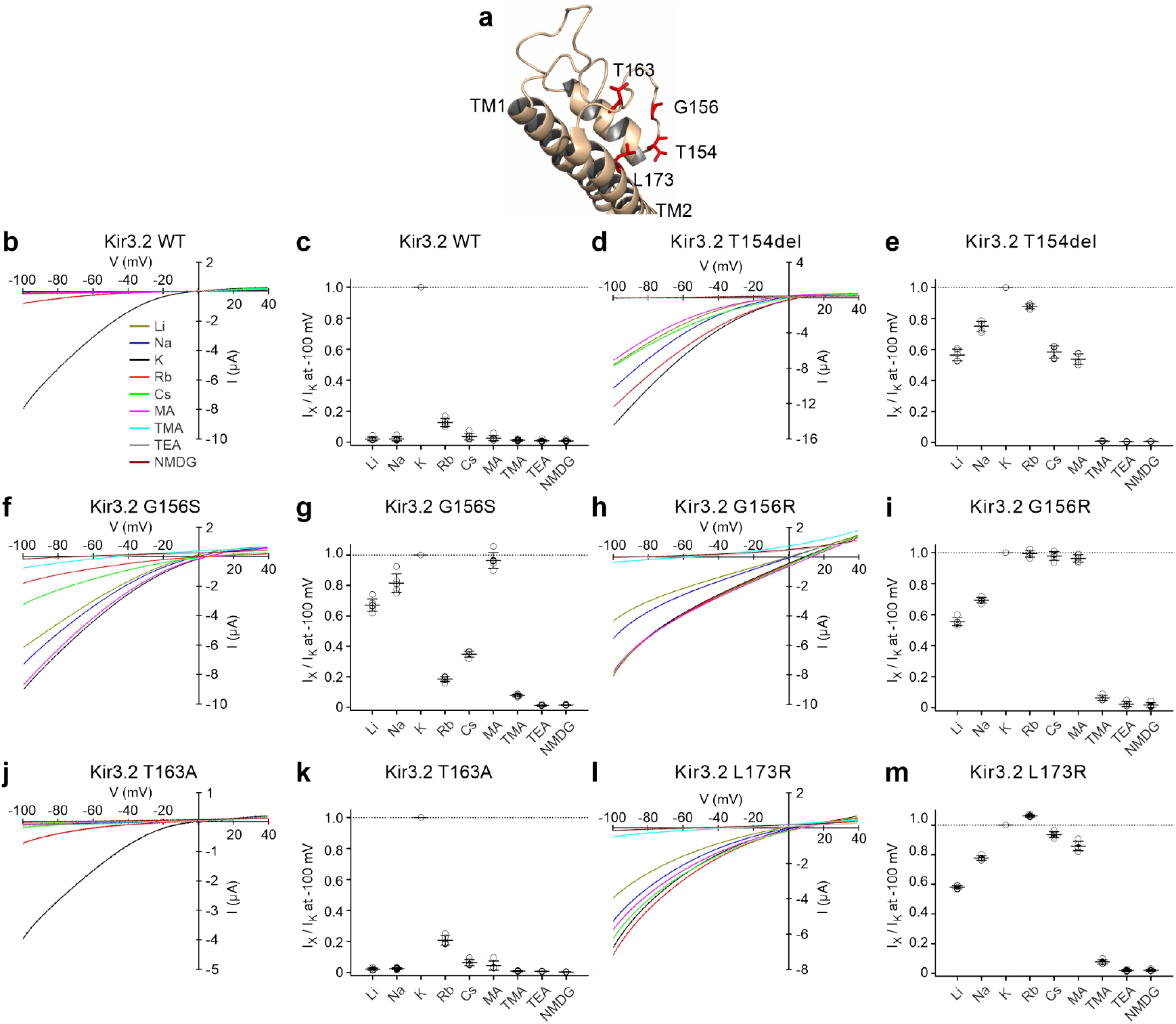
Permeation of various cations in inherited Kir3.2 mutants expressed in *Xenopus* oocytes. (**a**) The crystal structure of the pore-forming region of Kir3.2 WT (Protein Data Bank ID: 3SYA)(Whorton & MacKinnon, 2011). The positions of inherited mutations are shown and labelled in red in stick mode. The IV-relationship of Kir3.2 WT (**b**), T154del (**d**), G156S (**f**), G156R (**h**), T163A (**j**), and L173R (**l**) were recorded by a ramp pulse from −100 mV to +40 mV for 1 s in an extracellular solution containing 96 mM XCl, where the X is Li, Na, K, Rb, Cs, MA, TMA, TEA, or NMDG, respectively. (**c**, **e**, **g**, **i**, **k**, **m**) The current amplitude of X^+^ relative to K^+^ (I_X_/I_K_) of each mutant at −100 mV was calculated by normalization of I_K_ as 1. Data shown are individual values with means ± SD, n = 6 for WT and G156S; n = 5 for the other mutants.

By using a ramp voltage pulse from −100 mV to +40 mV, we observed that wild-type (WT) Kir3.2 showed a large K^+^ current and a minor Rb^+^ current at −100 mV (Figure 1b). Only negligible currents were observed when K^+^ was replaced with other cations. The current amplitude of each cation (I_X_) at −100 mV relative to that of K^+^ (I_K_) is referred to as I_X_ / I_K_ in Figure 1c. To analyze the ion selectivity quantitatively, the change of the reversal potentials by replacing the extracellular cations is usually used. In the case of GIRK channels which show strong rectification, however, the reversal potential is significantly affected by even a small leak current which hinders accurate analysis of P_X_ / P_K_. Isolation of the GIRK current component by using blockers was also not possible, due to the decreased sensitivity to pore blockers as shown in Figures 2, 3, 6 and 7. Thus, we did not perform the measurement of the reversal potential in this study except for those in Figure 3 h-j. We instead compared the current amplitude in various extracellular solutions. The information we present in this study to compare the ion selectivity is the profile of sequence of current amplitudes which reflects the extent of the permeation of various ions.

**Figure 2.**
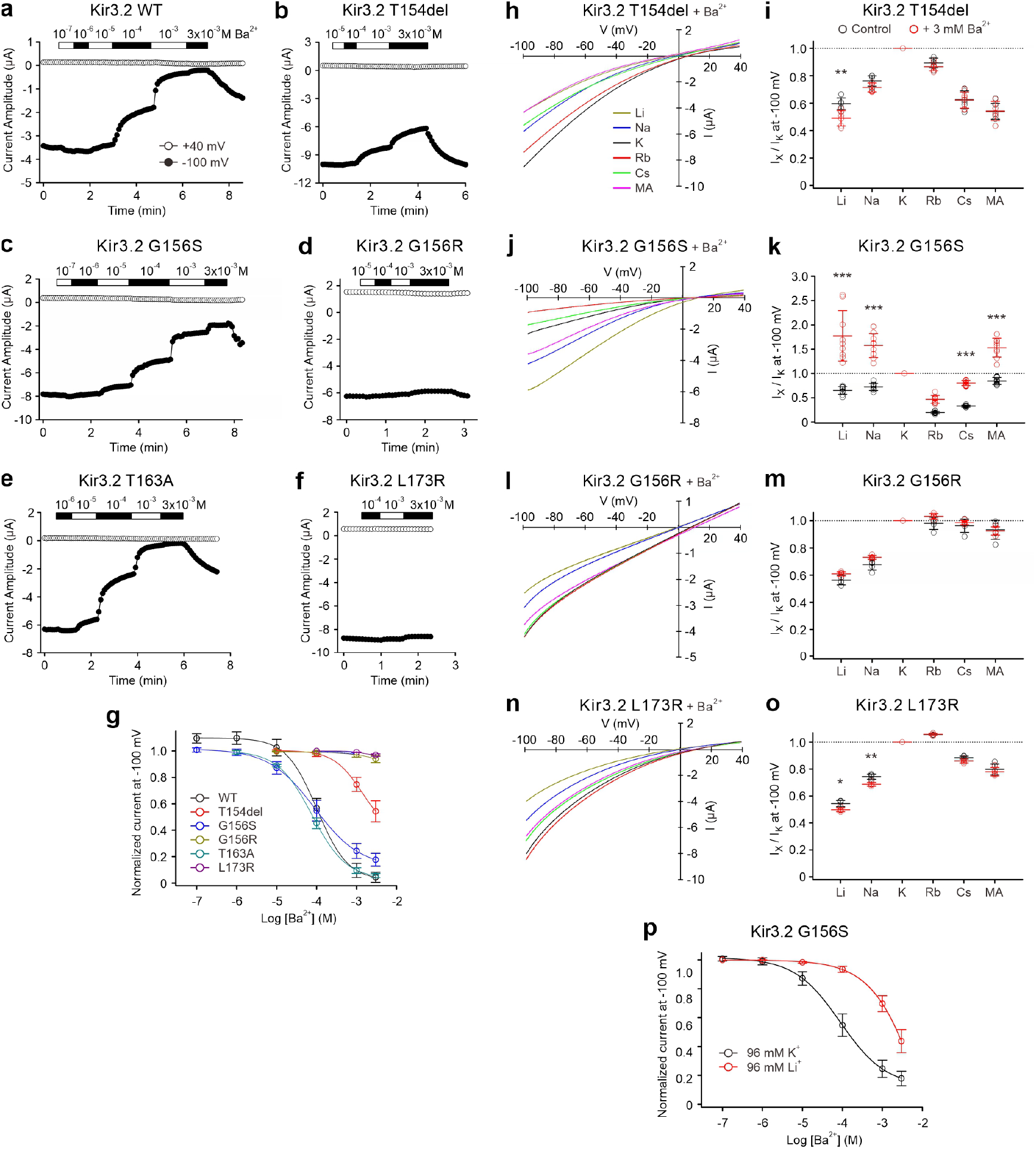
Effect of Ba^2+^ on the current of various cations in inherited Kir3.2 mutants expressed in *Xenopus* oocytes. (**a-f**) Representative time course of current change of Kir3.2 WT and mutants at −100 mV and +40 mV before and after applications of different concentrations of BaCl_2_ in the extracellular solution. (**g**) Dose-inhibition relationship at −100 mV. Data shown are means ± SD, n = 5 for each. The IV-relationship of Kir3.2 T154del (**h**), G156S (**j**), G156R (**l**), and L173R (**n**) were recorded by a ramp pulse from −100 mV to +40 mV for 1 s in an extracellular solution containing 96 mM XCl, where the X refers to Li, Na, K, Rb, Cs, or MA, respectively (**i**, **k**, **m**, **o**). The I_X_/I_K_ of each mutant at −100 mV before (black symbols) and after the application of 3 mM BaCl_2_ (red symbols). Data shown are individual values with means ± SD, n = 6 for T154del; n = 9 for G156S; n = 5 for the other mutants. * *P* < 0.05, ** *P* < 0.01, *** *P* < 0.001 depict the significance of statistical difference between the values in the absence and the presence of Ba^2+^. (**p**) Dose-inhibition relationship of Ba^2+^ on the K^+^ current (black symbols) or the Li^+^ current (red symbols) of Kir3.2 G156S, means ± SD, n = 5 for each.

**Figure 3.**
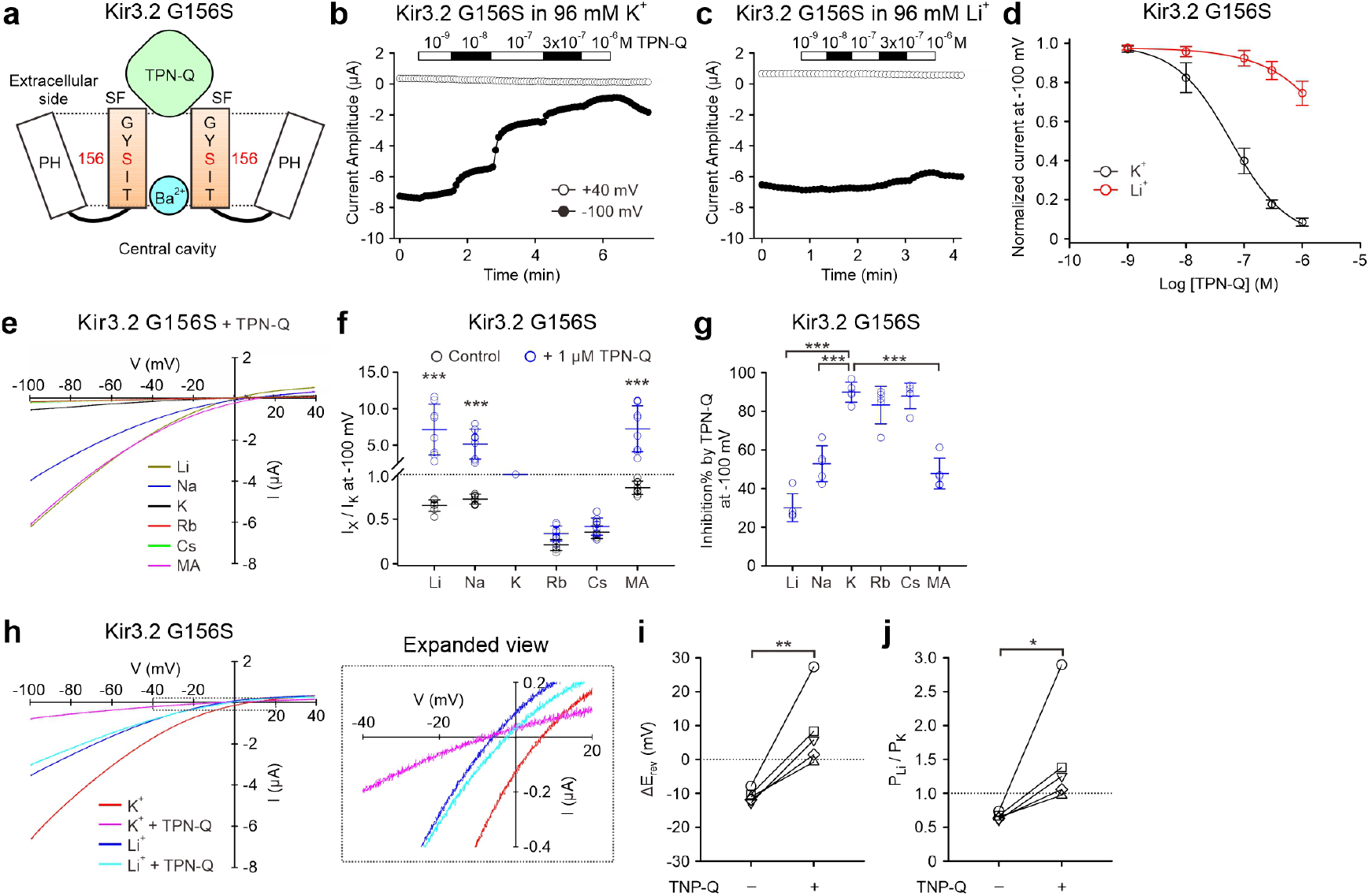
Effect of TPN-Q on the current of various cations of Kir3.2 G156S expressed in *Xenopus* oocytes. (**a**) A schematic drawing of the pore-forming region and the location of G156S of Kir3.2. Ba^2+^ and TPN-Q block the ion permeation route formed by the SF are drawn according to previous studies (Alagem et al., 2001; Doupnik et al., 2015; Hsieh et al., 2015; Patel et al., 2020). Time course of current change at −100 mV and +40 mV before and after applications of different concentrations of TPN-Q in the extracellular solution containing 96 mM KCl (**b**) or LiCl (**c**). (**d**) Dose-inhibition relationship of K^+^ current (black symbols) and Li^+^ current (red symbols) at −100 mV, means ± SD, n = 5 for each. (**e**) The IV-relationship of each cation in the presence of 1 μM TPN-Q. (**f**) The I_X_/I_K_ at −100 mV before (black symbols) and after the application of TPN-Q (blue symbols). Data shown are individual values with means ± SD, n = 8. *** *P* < 0.001 depict the statistical significance of difference between the absence and the presence of TPN-Q. (**g**) Inhibition percentage of cation currents at −100 mV by 1 μM TPN-Q, means ± SD, n = 5 for each, *** *P* < 0.001 as compared the K^+^ group. (**h**) The IV-relationship of K^+^ and Li^+^ current in the absence or presence of 1 μM TPN-Q. An expanded view of zero current range is shown in the right panel. (**i**) The shift of reversal potential when the extracellular Li^+^ was replaced with K^+^, means ± SD, n = 5. (**j**) The permeability of Li^+^ relative to the permeability of K^+^ (P_Li_/P_K_) were calculated from (**i**), means ± SD, n = 5. * *P* < 0.05, ** *P* < 0.01 depict the statistical significance of difference between the values in the absence and the presence of TPN-Q.

Deletion of Kir3.2 Thr154 in the SF resulted in inward currents induced by various extracellular cations in the order of K^+^ > Rb^+^ > Na^+^ > Li^+^ ≈ Cs^+^ ≈ MA^+^ (Figure 1d, e). Mutation of Kir3.2 Gly156 in the SF to Ser also showed abnormal inward currents but the sequence of current amplitudes produced by various cations was different from that of T154del (Figure 1f, g). Kir3.2 G156S showed larger Na^+^, Li^+^, MA^+^ currents than Rb^+^ or Cs^+^ currents. Since the homotetramers of the Kir3.4 SF mutant G151R showed very small currents from all extracellular cations, ion permeation could not be compared because of contamination from endogenous currents (Figure 1-figure supplement 1a). We, therefore introduced the mutation at the corresponding position of Kir3.2 (G156R) and observed that its ion selectivity was different from that of Kir3.2 G156S (Figure 1f-i). G156R showed a high selectivity to Rb^+^, Cs^+^ and MA^+^, and less inward rectification. The results show that mutation of Gly156 to different amino acid residues induces distinct conformational changes to the ion permeation pathway.

Kir3.4 T158A (T163A in Kir3.2; Figure 1a) in the extracellular loop after the SF also showed an abnormal ion selectivity (Figure 1-figure supplement 1b, c) which was similar to those of G156R and L173R in the TM2 of Kir3.2 (Figure 1a, h, i, l, m). A previous study suggested that this may be relevant to the eliminated hydrogen bonds of Thr158 with Pro128, Cys129 and Lys160, which are conserved among GIRK subtypes (Choi et al., 2011). However, when we introduced the corresponding mutation into Kir3.2 in the extracellular loop after the SF (T163A), it showed an intact ion selectivity and inward rectification like the WT (Figure 1b, c, j, k). This suggests that the hydrogen bonds formed between the threonine and the three conserved residues are not necessarily relevant to the ion selectivity change. Instead, the extracellular loop may play some roles to maintain the ion selectivity.

Essentially, no significant currents were observed in the WT and all mutants when replacing the extracellular cations with TMA^+^, TEA^+^ or NMDG^+^ (Figure 1 and Figure 1-figure supplement 1b, c), showing that these mutants did not allow the permeation of extra-large cations. These results demonstrate that most of the inherited mutations of Kir3.2 or Kir3.4 channels induce a conformational change with higher Rb^+^ and Cs^+^ permeability than for Na^+^ and Li^+^. However, a unique conformational change is induced by the G156S mutation of Kir3.2 with higher permeation of Na^+^ and Li^+^ than Rb^+^ and Cs^+^.

### Effect of Ba^2+^ on the ion selectivity of Kir3.2 G156S

To further characterize the property of ion permeation for each mutant, we examined the effect on ion selectivity of a well-established pore blocker of Kir channels, Ba^2+^. Ba^2+^ is known to block the ion permeation pore of Kir channels by strong interaction with the threonine residue located at the bottom of the SF (eg. Thr154 in Kir3.2) (Alagem, Dvir, & Reuveny, 2001; Hsieh, Kuo, & Huang, 2015). We first investigated the potency and efficacy of Ba^2+^ in the inhibition of K^+^ currents for WT and mutants of Kir3.2. Application of Ba^2+^ inhibited the K^+^ current of WT, G156S and T163A well (IC_50_: 105 ± 19 μM for WT; 95 ± 37 μM for G156S; 78 ± 11 μM for T163A, n = 5 for each), and inhibited T154del with an intermediate potency (Figure 2a-g). The sensitivity of G156R and L173R to the block by Ba^2+^ was very low. Kir3.4 T158A also showed a very low sensitivity to Ba^2+^ (Figure 2-figure supplement 1a, b).

We next examined the ion selectivity of mutants before and after the application of a high concentration of Ba^2+^ (3 mM), except Kir3.2 WT and the WT-like mutant T163A. For T154del, G156R, L173R of Kir3.2 or T158A of Kir3.4, application of Ba^2+^ did not influence the sequence of I_X_/I_K_ of cations tested (Figure 2h, i, l-o and Figure 2-figure supplement 1c, d). Unexpectedly, Ba^2+^ did change the sequence of ion selectivity of Kir3.2 G156S, which showed an apparent increase in the I_Li_/I_K_, I_Na_/I_K_ and I_MA_/I_K_ (Figure 2j, k). The current amplitude of Li^+^, Na^+^ or MA^+^ at −100 mV became larger than the K^+^ current in the presence of Ba^2+^. These results show that Kir3.2 G156S possesses a different ion permeation mechanism from that of the other mutants.

We examined the dose-inhibition relationship of Ba^2+^ on the K^+^ and the Li^+^ currents of Kir3.2 G156S (Figure 2p). The curve was shifted to the right when extracellular K^+^ was replaced with Li^+^, suggesting that Ba^2+^ blocks the Li^+^ permeation with a lower affinity than that of the K^+^ permeation. There are two possible explanations for the altered permeation and block of the G156S mutant as follows. (1) On top of the SF pathway blocked by Ba^2+^ with a high affinity, a novel ion permeation pathway is formed which allows permeation of Li^+^, Na^+^ and MA^+^ with a lower sensitivity to the block by Ba^2+^. (2) Ba^2+^ firmly binds to the threonine (Thr154) at the bottom of SF when K^+^ ions enter the SF, while the Ba^2+^ binding becomes unstable when Li^+^, Na^+^ or MA^+^ enter the SF pathway.

### Effect of TPN-Q on the ion selectivity of Kir3.2 G156S

To further clarify whether a novel ion permeation pathway besides the conventional SF route does exist in Kir3.2 G156S, we examined the effect of another pore blocker which binds to the GIRK channel at a different position than Ba^2+^. Here we used Tertiapin-Q (TPN-Q) known to specifically block Kir1.1 and GIRK channels by capping the SF pore from the extracellular side (Figure 3a) (Doupnik, Parra, & Guida, 2015; Patel, Kuyucak, & Doupnik, 2020). Application of TPN-Q inhibited the K^+^ current strongly and inhibited the Li^+^ current weakly in Kir3.2 G156S (Figure 3b-d). In the presence of TPN-Q, Li^+^, Na^+^ or MA^+^ current at −100 mV became approximately 5-10 times larger than the K^+^ current (Figure 3e, f). The I_Li_/I_K_, I_Na_/I_K_ and I_MA_/I_K_ were increased by the application of TPN-Q, and this is consistent with the effect of Ba^2+^ application (Figure 3f and 2k). We calculated the percent inhibition of the current for each cation and found that 1 μM of TPN-Q inhibited about 83~90% of the K^+^, Rb^+^, or Cs^+^ currents, 52.9 ± 9.3 % of the Na^+^ current, 47.8 ± 7.9 % of the MA^+^ current, and 30.1 ± 7.2 % of the Li^+^ current (n = 5 for each, Figure 3g). One possible interpretation of the results would be as follows: K^+^, Rb^+^, or Cs^+^ predominantly permeate through the channel via the SF pathway, with a very a limited permeation via the novel route; in the case of Na^+^ or MA^+^, half and half permeation occurs through the two pathways; Li^+^ permeates predominantly through the novel route which is less sensitive to the block by TPN-Q.

To clarify whether a novel permeation pathway with a high Li^+^-permeability does exist in Kir3.2 G156S in addition to the SF route with a high K^+^-permeability, we examined the permeability of Li^+^ (P_Li_) relative to the permeability of K^+^ (P_K_) by measuring the shift of reversal potentials (ΔE_rev_) upon replacement of the extracellular Li^+^ with the same concentration of K^+^ (Figure 3h-j). In the absence of TPN-Q, E_rev_ of Li^+^ current was −2.7 ± 5.8 mV (n = 5) and was shifted to the right in K+ solution (8.1 ± 5.4 mV, n = 5), showing that P_K_ is larger than P_Li_. E_rev_ of the K^+^ current shifted to the left on application of 1 μM TPN-Q (−9.6 ± 11.1 mV, n = 5) and shifted to the right upon replacement of K^+^ with Li^+^ (−1.2 ± 6.3 mV, n = 5), showing that P_K_ was decreased by TPN-Q and that P_K_ is smaller than P_Li_ in the presence of TPN-Q. Taken together, when the SF pathway was blocked by TPN-Q, a high P_Li_/P_K_ was observed presumably due to the novel ion permeation route. When the SF route is not blocked, a lower P_Li_/P_K_ was observed as a result of the presence of two pathways.

### Activation effect of G_βγ_ and Ivermectin on Kir3.2 G156S

Previous studies showed that heteromeric channels comprised of Kir3.2 G156S and Kir3.1 are insensitive to G_βγ_-stimulation (Navarro et al., 1996). Here we examined whether homomeric channels of Kir3.2 G156S are also insensitive to the G_βγ_-stimulation by coexpressing Kir3.2 G156S with the muscarinic M2 receptor (M2R). We observed that an application of 1 μM ACh increases the K^+^ current of Kir3.2 WT but does not apparently increase that of G156S (Figure 4), showing that homotetramers of Kir3.2 G156S are also insensitive to G_βγ_-stimulation as compared with the WT. We also examined the effect of another activator of Kir3.2 channels, Ivermectin (IVM), which directly interacts with the channel as we reported previously (Chen & Kubo, 2018; Chen, Tateyama, Fukata, Uesugi, & Kubo, 2017). Applications of 10 μM IVM induced the current increase in both the Kir3.2 WT and G156S mutant. This shows that the limited response of the G156S mutant to ACh stimulation is not due to the high basal activity reaching a saturation level, since IVM can further increase the current amplitude. Taken together, the G156S mutation may induce a conformational change that occurs not only in the SF region but also in the structural components which play roles in the G_βγ_-coupling-channel activation linkage.

**Figure 4.**
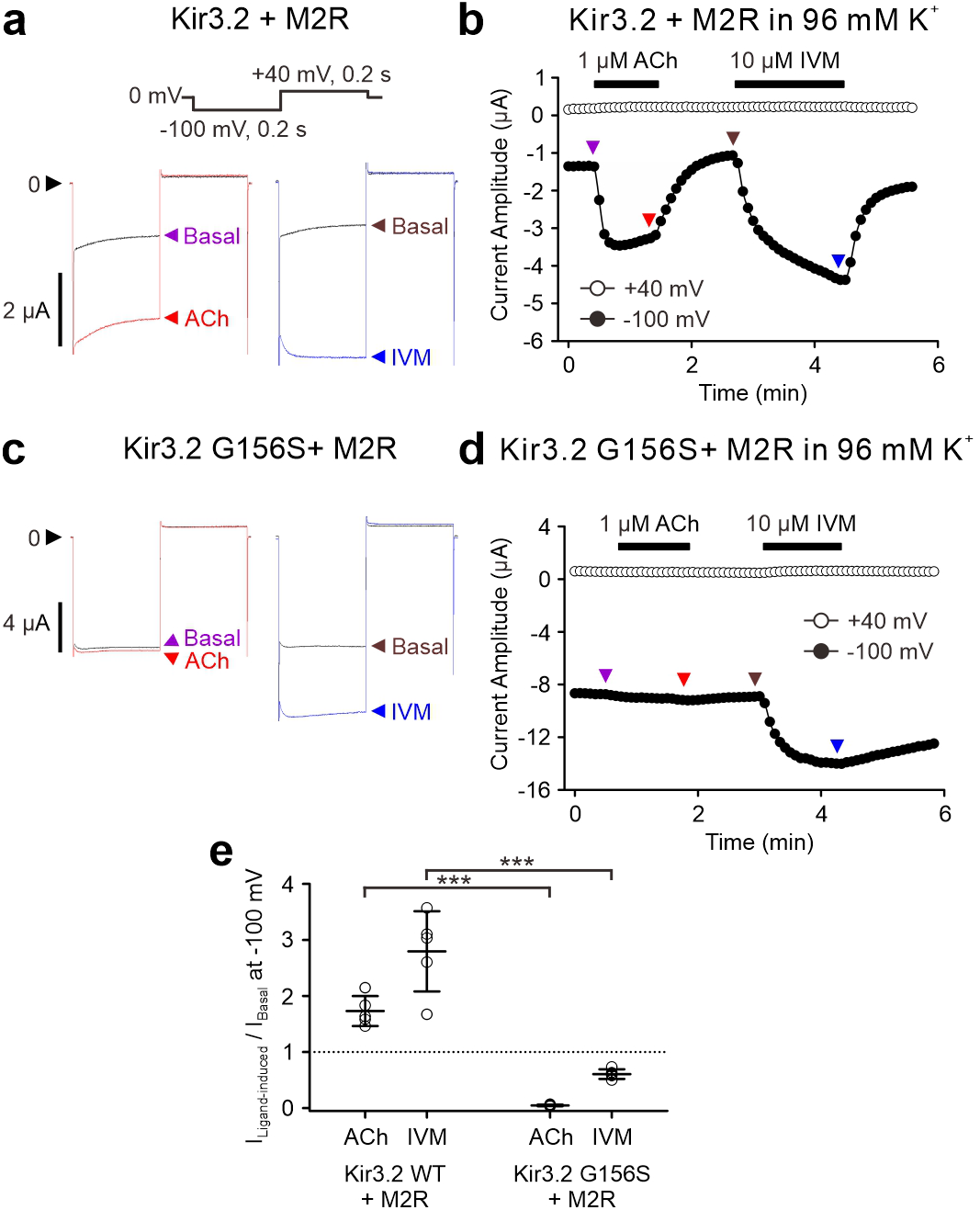
Effects of activators of GIRK channels on the current of Kir3.2 G156S expressed in *Xenopus* oocytes. Current of oocytes expressing Kir3.2 WT with M2R (**a**, **b**) or Kir3.2 G156S with M2R (**c**, **d**) were recorded in an extracellular solution containing 96 mM K^+^ with a protocol shown above. Current traces before or after the application of 1 μM ACh or 10 μM Ivermectin (IVM) at the time indicated by arrows (**b**, **d**) are shown in (**a**, **c**). (**e**) The amplitudes of ACh-induced current or IVM-induced current (I_Ligand-induced_) relative to the basal currents (I_Basal_) were calculated by normalization of I_Basal_ as 1. Data shown are individual values with means ± SD, n = 5 for each. *** *P* < 0.001 indicates the statistical significance of the difference between the WT and the mutant.

### Single-channel recordings of Kir3.2 G156S show two types of ion conducting events

We speculated that there should be two types of ion conducting events in G156S, one generated by K^+^ permeation through the SF route and the other generated by Li^+^ permeation through the novel route. To verify this hypothesis, we performed cell-attached single-channel recordings using mouse fibroblasts heterologously expressing Kir3.2 G156S with a pipette solution of 140 mM K^+^, 140 mM Li^+^, or a mixture of 70 mM K^+^ and 70 mM Li^+^, in the absence or presence of TPN-Q respectively (Figure 5). In 140 mM K^+^ at −100 mV, we observed inward currents with amplitudes up to ~2 pA (Figure 5a). When 400 nM TPN-Q was added to the pipette solution, these events with a relatively large current amplitude were largely absent and replaced by infrequently observed events with a very small current amplitude (< 0.2 pA) (Figure 5b). Occasional amplifier resets were unavoidable during these long recordings due to the need to reset the headstage output periodically as part of the capacitive feedback system used to achieve low noise recordings (Figure 5b). The results demonstrate that the major events of larger conductance by K^+^ influx of Kir3.2 G156S were blocked by TPN-Q (Figure 5g).

**Figure 5.**
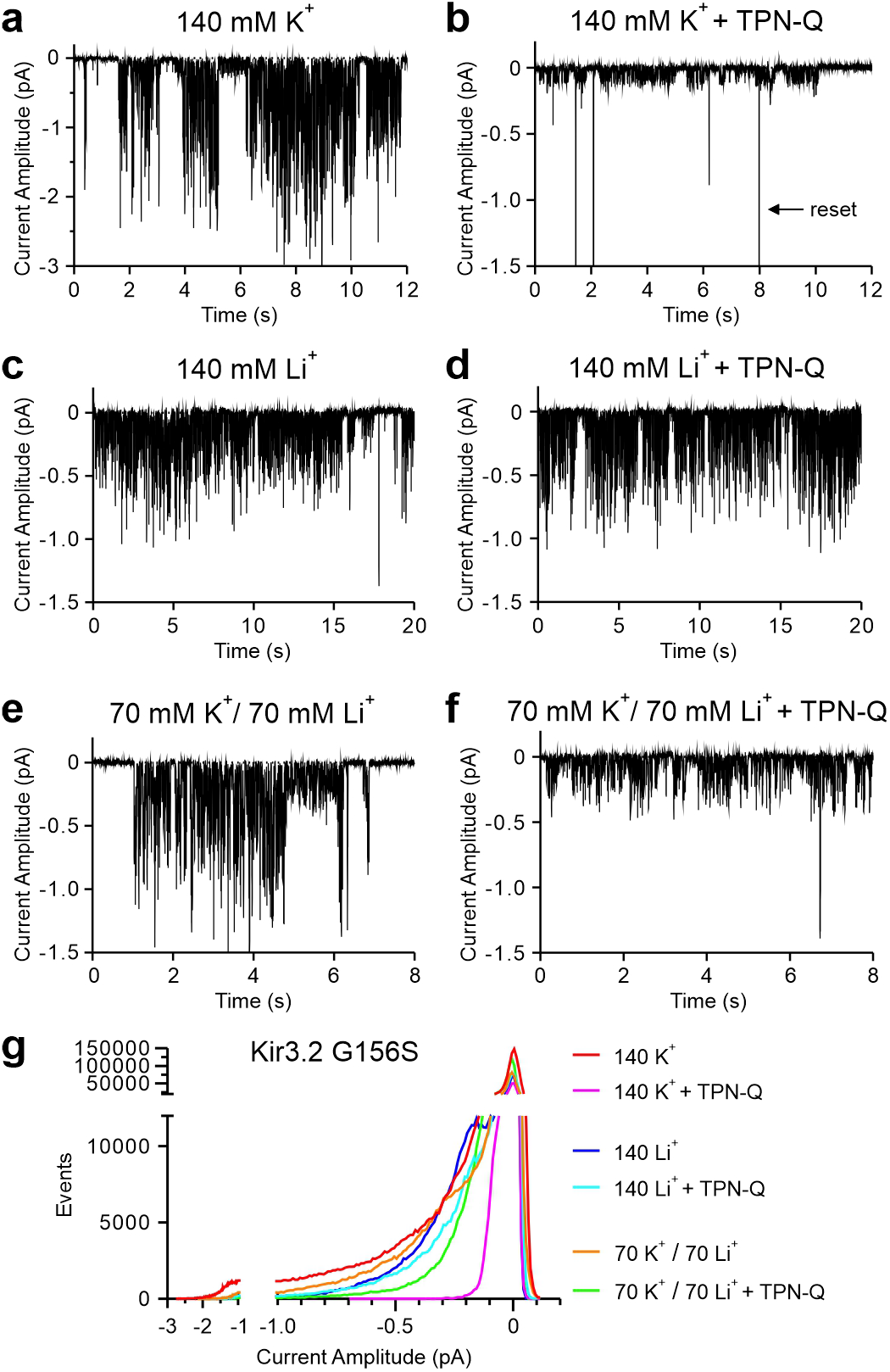
Single-channel activity of Kir3.2 G156S mutant expressed in mouse fibroblasts. Representative single channel recordings of the Kir3.2 G156S mutant at −100 mV with 140 mM K^+^ (**a**, **b**), 140 mM Li^+^ (**c**, **d**) or a mixture or 70 mM K^+^/70 mM Li^+^ (**e**, **f**) in the recording pipette, with or without 400 nM TPN-Q as indicated. Occasional artifacts due to amplifier resets are unavoidable and an example is indicated in (**b**). (**g**) Current amplitude histograms of net ion conduction events in the absence or presence of TPN-Q with three types of solution compositions. Data are pooled from n = 3 to 5 for each condition.

In 140 mM Li^+^, inward current events with a peak amplitude smaller than 1 pA were recorded (Figure 5c), smaller than those observed in 140 K^+^ in Figure. 5a. Inclusion of 400 nM TPN-Q in the pipette did not apparently change the activity and the amplitude of Li^+^ current (Figure 5c, d), showing that the major events caused by Li^+^ influx through Kir3.2 G156S were not blocked by TPN-Q (Figure 5g).

To clarify whether the K^+^ conducting events and Li^+^ conducting events are present in the same recording, we performed recordings with 70 mM K^+^ and 70 mM Li^+^ solution in the pipette. In the current recordings with a slow ramp pulse from +30 mV to −150 mV for 40 s, the conductance was reduced by the presence of 400 nM TPN-Q in the pipette, at the tested membrane potentials (Figure 5-figure supplement 1). In the current recordings held at −100 mV in the absence of TPN-Q, two types of inward current events were detected; one type of event with a peak amplitude > 0.5 pA and another type of event with a peak amplitude < 0.5 pA (Figure 5e). The events with a larger amplitude between 4 s and 5 s in Figure. 5e were interpreted to be the sum of simultaneous activities of two type of events. In the presence of 400 nM TPN-Q, only the smaller events with < 0.5 pA remained (Figure 5f), and similar events were also observed in Figure. 5e between 5 s and 6 s. These data together with those in Figure 5a-d suggest that the single channel events with a larger amplitude blocked by TPN-Q involve K^+^ influx, and the events with a smaller amplitude which remained in the presence of TPN-Q involve Li^+^ influx. These results demonstrate that two types of ion conducting events are present in the same recording of Kir3.2 G156S. This supports that a novel ion permeation pathway in addition to the SF route is formed by the G156S mutation in the Kir3.2 channel.

### Mutations in the pore helix located behind the SF also form a Li^+^-permeable pathway

To reveal the location of the novel ion permeation route, we introduced a point mutation into the PH located behind the SF, since there is a pocket accessible from the extracellular side surrounded by the side chains of amino acid residues of the SF and PH in the WT. We previously reported that the S148F mutant located at the center of the PH in Kir3.2 is Na^+^-permeable (Chen et al., 2019) (Figure 6a). Here we examined the ion permeation of various cations and observed that this mutant permeates Li^+^, Na^+^, Rb^+^, Cs^+^ and MA^+^ and their I_X_/I_K_ are 0.92 ± 0.04 for MA^+^ and about 0.43~0.60 for the other cations (n = 6 for each, Figure 6b, d). The dose-inhibition relationship for Kir inhibitors on Kir3.2 S148F showed that 3 mM Ba^2+^ blocks 72.8 ± 3.9 % of the K^+^ current and 1 μM TPN-Q blocks 48.0 ± 16.0 % (n = 5 for each, Figure 6c). In the presence of Ba^2+^ or TPN-Q, the I_Li_/I_K_, I_Na_/I_K_, and I_MA_/I_K_ became larger than those in the absence of these inhibitors (Figure 6d), demonstrating that a novel route, which is permeable to Li^+^, Na^+^ and MA^+^ and insensitive to the block by TPN-Q, exists also in the S148F mutant.

**Figure 6.**
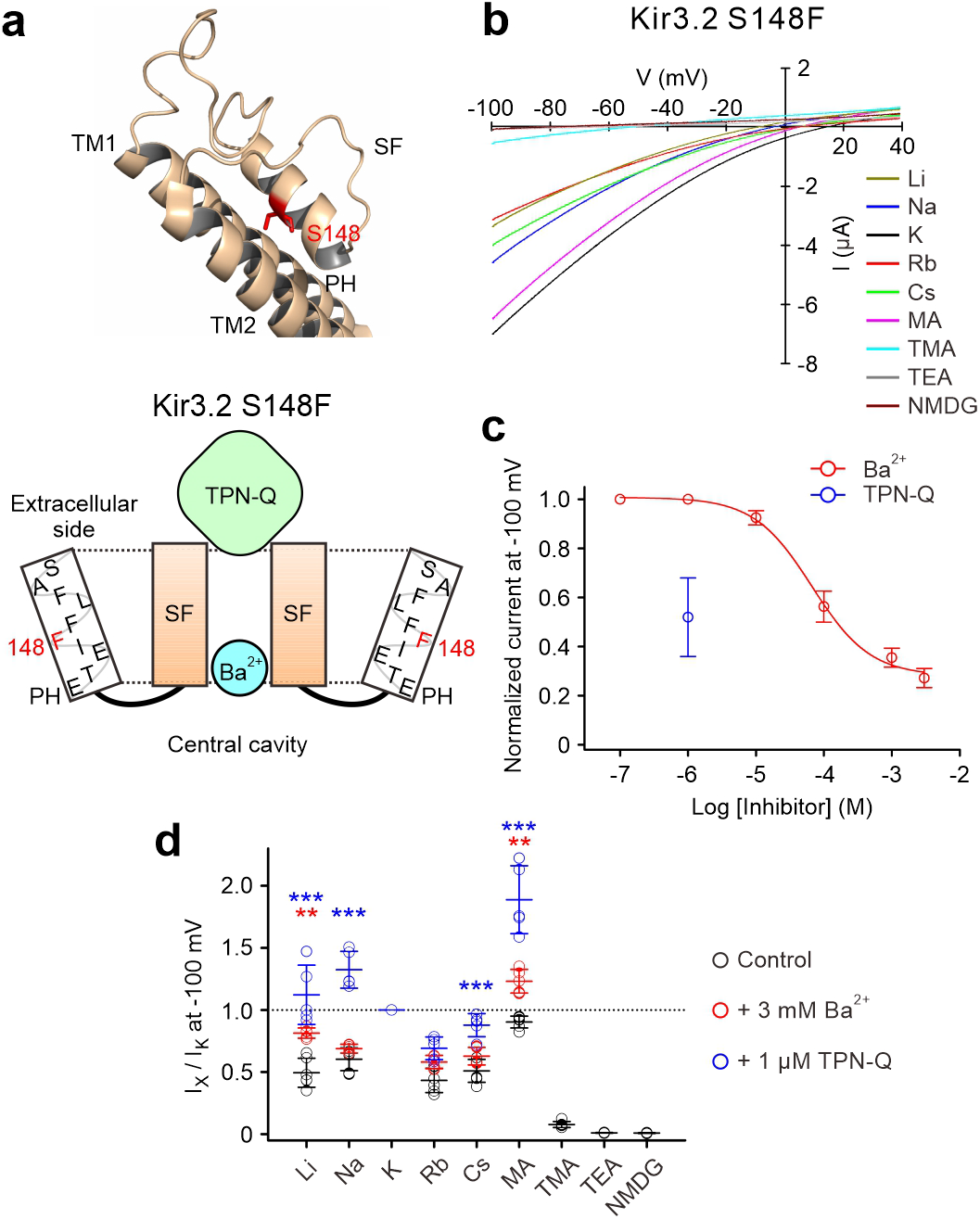
Permeation of various cations and the effect of blockers on the currents of Kir3.2 S148F expressed in *Xenopus* oocytes. (**a**) The position of S148 (labelled in red) at the pore helix (PH) of Kir3.2 WT. (**b**) The IV-relationship of each cation. (**c**) Dose-inhibition relationship of Ba^2+^ (red symbols) and TPN-Q (blue symbols) on K^+^ current at −100 mV, means ± SD, n = 5 for each. (**d**) The I_X_/I_K_ at −100 mV in the absence of blocker (black symbols), and in the presence of 3 mM Ba^2+^ (red symbols) or 1 μM TPN-Q (blue symbols). Data shown are individual values with means ± SD, n = 6 for control group; n = 5 for the other groups. ** *P* < 0.01, *** *P* < 0.001 depict the statistical significance of difference between the absence and the presence of Ba^2+^ (red *) or TPN-Q (blue *).

Unexpectedly, mutation of Thr151, which is located at the bottom of the PH (Figure 7a), to Ala induced an extremely high rate of Li^+^ permeation, even in the absence of SF route blockers (Figure 7b). The K^+^ current of Kir3.2 T151A was very small and the Li^+^, Na^+^ and MA^+^ currents were far larger than the K^+^ current (I_Li_/I_K_: 19.1 ± 9.2; I_Na_/I_K_: 11.8 ± 6.1; I_MA_/I_K_: 17.7 ± 7.2, n = 10 for each) (Figure 7b, d). The T151C mutation also showed high I_Na_/I_K_ (I_Na_ > I_K_). However, only a limited number of successful recordings were obtained, due to the very unhealthy condition of the oocytes expressing the T151C mutant (Figure 7-figure supplement 1). Mutation of Thr151 to Ser retained intact K^+^ selectivity as WT, while mutation of Thr151 to Asp, Asn, Lys, Tyr, or Phe triggered permeation of Na^+^, but I_Na_ was still smaller than I_K_ (Figure 7-figure supplement 1b). Oocytes expressing other T151 mutants showed very small currents (Figure 7-figure supplement 1a). These results show that the Ala mutant at this position increases the Na^+^, Li^+^ permeation. Since the K^+^ current of the T151A mutant was very tiny, hindering detailed dose-inhibition relationship analysis, we compared the inhibiting effect of blockers on the Li^+^ current (Figure 7c). Application of 3 mM Ba^2+^ blocked 59.5 ± 6.0 % (n = 5) of the Li^+^ current and 1 μM TPN-Q only slightly reduced the Li^+^ current (10.1 ± 2.5 %, n = 5, Figure 7c). The sequence of I_X_/I_K_ did not change apparently by applying blockers (Figure 7d), suggesting that T151A mutant has only a novel ion permeation pathway. The results demonstrate that this ion permeation route allows Li^+^, Na^+^ and MA^+^ permeation better than K^+^ or Rb^+^, and is insensitive to the block by TPN-Q. These are consistent with the features of the novel pathway identified in Kir3.2 G156S. It appears that the T151A mutation induces a conformational change which forms a Li^+^-permeable pathway in the pocket surrounded by the SF and PH residues, that the SF route is collapsed to inhibit K^+^ permeation, and that Ba^2+^ enters the novel ion permeation pathway and inhibits Li^+^ current with a lower binding affinity than that for the SF route in WT (Figure 7a). This is similar to the dose-inhibition curve for Ba^2+^ on the Li^+^ currents of Kir3.2 G156S, that is shifted to the right compared to that of the K^+^ current (Figure 2p).

**Figure 7.**
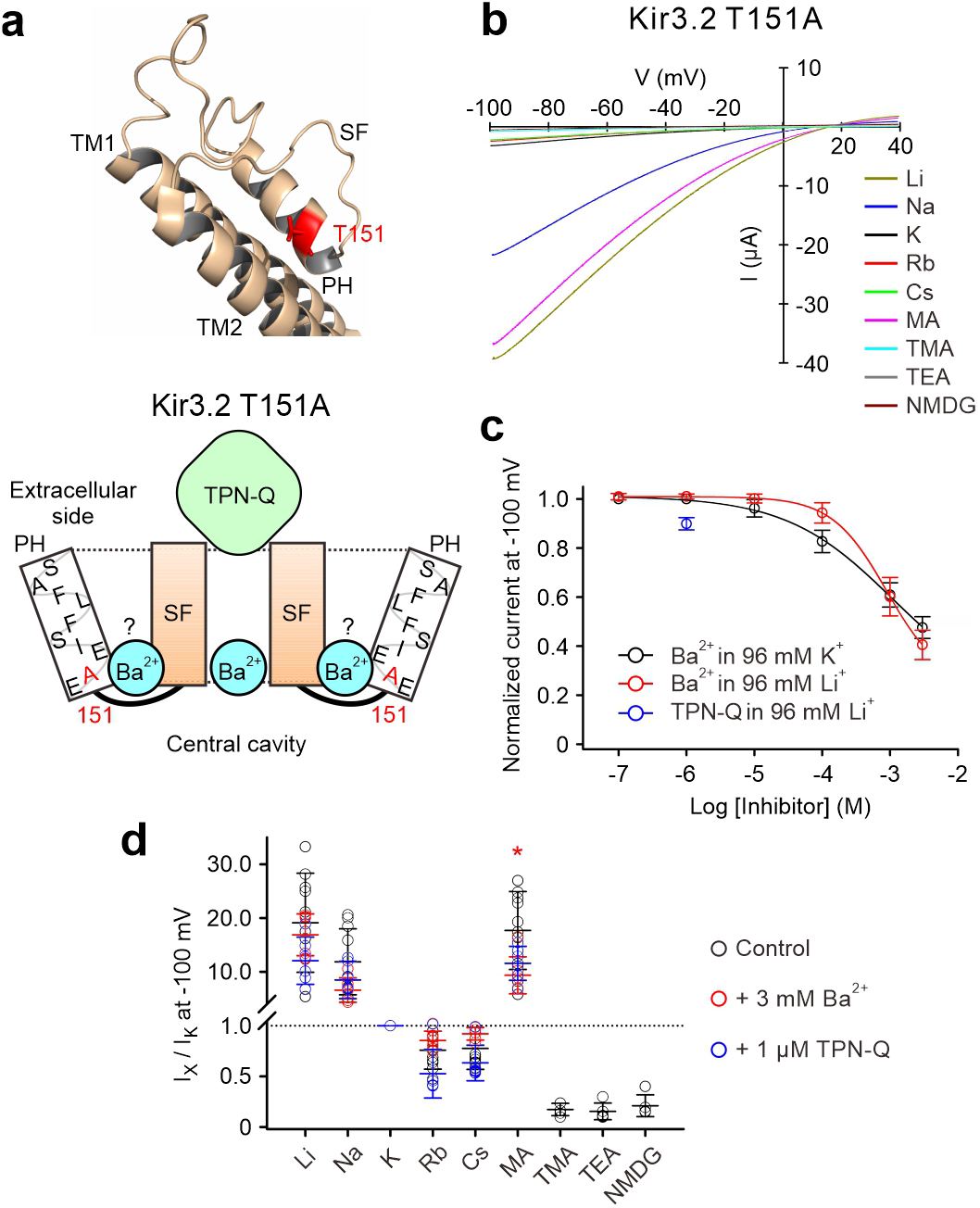
Permeation of various cations and the effect of blockers on the currents of Kir3.2 T151A expressed in *Xenopus* oocytes. (**a**) The position of T151A (labelled in red) in the pore helix of Kir3.2 WT. (**b**) The IV-relationship of each cation. (**c**) Dose-inhibition relationship of Ba^2+^ on K^+^ current (black symbols) or Li^+^ current (red symbols), and TPN-Q on Li^+^ current (blue symbols) at −100 mV, means ± SD, n = 5 for each. (**d**) The I_X_/I_K_ at −100 mV in the absence of blocker (black symbols), and in the presence of 3 mM Ba^2+^ (red symbols) or 1 μM TPN-Q (blue symbols). Data shown are individual values with means ± SD, n = 5 to 10 for control group; n = 7 for Ba^2+^ group; n = 6 for TPN-Q group. Red * *P* < 0.05 depict the statistical significance of the difference between the absence and the presence of Ba^2+^. There were no significant differences between the absence and presence of TPN-Q are shown.

During the recordings, we observed that the T151A mutant shows a very slow current increase or decrease when exchanging extracellular K^+^ to Na^+^ or Na^+^ to K^+^ as compared with the Kir3.2 WT (Figure 7-figure supplement 2). However, when we exchanged the extracellular Na^+^ to the impermeable NMDG^+^ or NMDG^+^ to Na^+^, the current change kinetics of T151A were as fast as those in WT. This suggests that the cations, which occupy the novel pathway, might strongly interact with the pathway lining residues.

## Discussion

We have presented the following in this study: (1) Kir3.2 G156S permeates Li^+^ and Na^+^ better than Rb^+^ and Cs^+^, while the other inherited mutants of GIRK channels permeate Rb^+^ and Cs^+^ better than Li^+^ and Na^+^ (Figure 1); (2) applications of SF pore blockers increase the Li^+^ and Na^+^-selectivity of Kir3.2 G156S but do not alter those of the other mutants (Figures 2 and 3); (3) single-channel recordings of Kir3.2 G156S show events of K^+^ influx which are blocked by extracellular TPN-Q, and also those of Li^+^ influx which are not blocked by TPN-Q (Figure 5); (4) Kir3.2 S148F also shows an increased Li^+^-selectivity in the presence of pore blockers (Figure 6); (5) Kir3.2 T151A shows an extremely high Li^+^-selectivity which is not influenced by the application of pore blockers (Figure 7).

Our results demonstrate that a novel ion permeation pathway allowing Li^+^, Na^+^, or MA^+^ influx is formed in Kir3.2 G156S, besides the conventional central pore route formed by the SF which is blocked by Ba^2+^ or TPN-Q (Figures 2k and 3f). This novel ion permeation route is similar to a side entry pathway for Na^+^ permeation through the sodium potassium (NaK) channel (Roy et al., 2021; Shi et al., 2018). The SF of the NaK channel is constructed by TVGDG, with D66 diverging from the conserved SF sequence of K^+^ channels. NaK channels permeate K^+^ in a symmetric conformation of the SF, while another asymmetric conformation of the lower part of the SF, composed of different subunits, is necessary for Na^+^ conduction (Shi et al., 2018). When D66 of the SF flips, Na^+^ associates with the side entry binding site behind the SF pore (Roy et al., 2021). K^+^ was also shown to associate with the side entry binding site, in addition to the canonical K^+^ binding sites within the SF (Roy et al., 2021). In our single-channel recordings of G156S, K^+^ conducting events with a tiny current amplitude were infrequently observed in the presence of TPN-Q (Figure 5b), suggesting that K^+^ may also slightly permeate through the novel ion permeation route of Kir3.2 G156S. Mutants of the voltage-gated K^+^ channel, Kv1.2 V370C or Kv1.3 V388C, are also suggested to possess an unconventional cation permeation pathway, named the σ-pore behind the SF pore, which permeates cations in a sequence of efficacy of Li^+^ > Na^+^ > NH ^+^ > Cs^+^ > K^+^ > Rb^+^ (Prutting & Grissmer, 2011; Tyutyaev & Grissmer, 2017, 2018). The reported Li^+^-preferable permeation of the σ-pore is consistent with the feature of the novel ion permeation pathway which we have identified in Kir3.2 G156S.

The G156S mutation of mouse Kir3.2 causes the loss of cerebellar granule cells in *weaver* mice (Patil et al., 1995). Blocking of Na^+^ flux through *weaver* Kir3.2 channels by MK-801 and QX-314 at the early stage of development is known to rescue the survival and differentiation of *weave*r granule neurons (Kofuji et al., 1996). The inherited mutations of human Kir3.4, G151R and L168R which correspond to Gly156 in the SF and Leu173 in the TM2 of mouse Kir3.2, are reported to increase aldosterone synthesis (Choi et al., 2011; Velarde-Miranda, Gomez-Sanchez, & Gomez-Sanchez, 2013). Macrolides inhibits these mutants but not the WT channel, and potentially serves as a diagnostic tool for APA patients harboring these mutations (Maiolino et al., 2018; Scholl et al., 2017). We observed that the Kir3.2 G156R mutant shows a Rb^+^, Cs^+^-preferable permeation which is insensitive to block by Ba^2+^ (Figure 2). This is different from the features of Kir3.2 G156S, which is Na^+^, Li^+^-preferable and Ba^2+^-sensitive, suggesting that mutations of Gly156 to different amino acid residues induced distinct conformational changes in the permeation pathway. Kir3.2 L173R also showed Rb^+^, Cs^+^-preferable and Ba^2+^-insensitive features like that of Kir3.2 G156R. This may be due to the dilated SF in these mutants that underlies the nonselective conformation of the SF with reduced sensitivity to the Ba^2+^ block, as suggested in a study of KirBac1.1 using single-molecule FRET analysis (S. Wang et al., 2019).

The Ba^2+^ blockade in a Ca^2+^-activated K^+^ channel, MthK, is known to be modulated by extracellular cations (Guo, Zeng, Cui, Chen, & Ye, 2014). There are four hydrated K^+^-binding sites (S1~S4) in the SF of K^+^ channels (Doyle et al., 1998; Zhou, Morais-Cabral, Kaufman, & MacKinnon, 2001). When the MthK channel pore is filled with K^+^, Ba^2+^ blocks ion permeation by staying at the S3 and S4 sites predominantly, while Ba^2+^ stays at the S2 site in a Na^+^ environment, which weakens the block (Guo et al., 2014). Therefore, it is possible that the Ba^2+^ block of Kir3.2 G156S in different extracellular cation environments could be different, resulting in the increase of I_X_/I_K_ in the presence of Ba^2+^ (Figure 2). However, this possibility is unlikely, because another inhibitor TPN-Q, which blocks the pore without entering the SF, but by capping the pore instead, also shows similar increases of I_X_/I_K_ for the same type of cations in Kir3.2 G156S (Figures 2k and 3f). These results suggest that the alteration of the ion selectivity by application of blockers is unlikely to be due to the modulation of blocker binding in the SF pore by extracellular cations. A similar phenomenon was also observed in Kir3.2 S148F (Figure 6), further supporting that a Li^+^-permeable pathway which is not blocked by Ba^2+^ or TPN-Q, exists in Kir3.2 G156S and S148F mutants. We previously introduced various amino acid residues at the S148 position in Kir3.2 and found that a nonpolar and bulky side chain at this position make the channel permeable to Na^+^ (Chen et al., 2019). We also demonstrated that T151A in the PH contributes to the formation of the Na^+^, Li^+^-conducting route behind the SF (Figure 7), suggesting important roles of PH residues in the modulation of ion selectivity.

Based on the sequence of permeation for various ions and the presence or absence of sequence change by application of blockers, it was shown that G156S (Figures 2k and 3f) and S148F (Figure 6d) of Kir3.2 both have the modified SF route which permeates Rb^+^ and Cs^+^ well and the novel pathway which permeates Li^+^ and Na^+^ preferably. It was also shown that T151A has only the novel pathway (Figure 7d), whereas T154del (Figure 2i), G156R (Figure 2m) and L173R (Figure 2o) of Kir3.2 have only the modified SF pathway. The sequence of ion preference of the G156R and L173R mutants (Rb^+^, Cs^+^-preferable permeation) fits to I – IV of the Eisenman sequence, suggesting that the modified SF pathway is a weak-field-strength site type, where the site-interaction energy is less than the hydration energy (Eisenman, 1962; Hille, 2001). In the case of the G156S mutant, the sequence of ion preference in the absence of blockers does not fit to any of the Eisenman sequences, further supporting the current is a mixture via multiple pathways. However, G156S in the presence of blockers or T151A in the absence/presence of blockers, show ion preferences which fit to XI or X of the Eisenman sequences, suggesting that the novel ion permeation pathway is a strong-field-strength site type, where the site-interaction energy overwhelms the hydration energy (Eisenman, 1962; Hille, 2001). The slow kinetics of current amplitude change of T151A upon the exchange of extracellular cations also supports the strong interaction of ions with their binding sites (Figure 7-figure supplement 2). It is likely that the novel pathway is located in the pocket between the PH and side chain of the SF, behind the central SF, and further studies are needed to elucidate the detail of the whole route.

We observed in the single channel recording analysis of Kir3.2 G156S in K^+^ and Li^+^ that there are two types of events, a larger one presumably carried by K^+^, sensitive to the block by TPN-Q and a smaller one, presumably carried by Li^+^, resistant to the TPN-Q block (Figure 5e, f). If permeation through both pathways is independent and allow permeation simultaneously, it is naturally expected the sum height of two types of single channel events would be observed. At 4 s - 5 s of the recording in Figure 5e, larger events were observed, which could be an indication of overlapping activities of both types. Further detailed analysis was not performed due to the difficulty of the precise estimation of the small current amplitudes of fast gating events. It may be natural to expect that there will be four novel side pathways behind the SF in one tetrameric channel. If this is the case, multiple levels of single channel current amplitudes might be observed. However, due to the relatively low open probability, the multiple levels of overlapped activities could not be clearly observed. Thus, a conclusion as to the number of novel pathways in one tetramer could not be drawn.

In conclusion, our results reveal the presence of a novel ion permeation pathway besides the SF route in Kir3.2 G156S. This provides us with novel insight into the mechanisms underlying the loss of K^+^ selectivity induced by inherited mutations in GIRK channels. The design of drugs that selectively block the novel cation permeation route alone to eliminate the abnormal cation fluxes may have the potential to develop a novel treatment for Kir disorders such as for KPLBS patients with the Kir3.2 G154S mutation (corresponding to mouse G156S) at an early stage of life.

## Materials and Methods

### Ethical approval of animal experiments

All animal experiments in this study were approved by the Animal Care Committee of the National Institutes of Natural Sciences (an umbrella institution of National Institute for Physiological Sciences, Japan), and were performed in accordance with its guidelines.

### Mutagenesis and cRNA preparations

We constructed mutants of Kir3.2 and Kir3.4 and prepared their cRNA as described previously (Chen et al., 2019). By PCR using PfuUltra II Fusion HS DNA Polymerase kit (Agilent technologies, Santa Clara, CA, USA) and the primers (Supplementary Table 1), we introduced point mutations in mouse Kir3.2 (GenBank accession no.: AF040051) or rat Kir3.4 (GenBank no.: L35771) which were confirmed by DNA sequencing. Their cDNA were linearized by restriction enzymes, and the complementary RNA were transcribed by the mMessage T3 or T7 mMachine Kit (Ambion, Austin, TX, USA). The cRNA of Kir3.2 (25 ng oocyte^−1^ for the wild-type, T154del, G156R or T163A; 5 ng oocyte^−1^ for L173R or T151A; 0.5 ng oocyte^−1^ for G156S or S148F), Kir3.4 (25 ng oocyte^−1^ for the G151R or T158A), and porcine muscarinic M2 receptor (M2R, GenBank no.: X04708, 5 ng oocyte^−1^) were injected into *Xenopus* oocytes.

### Oocyte preparations

We purchased adult female *Xenopus laevis* from Hamamatsu Seibutsu Kyouzai (Hamamatsu, Japan). Isolation of oocytes from the frogs were performed as described previously (Chen et al., 2019). Frogs were anaesthetized with 0.15% tricaine (Sigma-Aldrich, St. Louis, MO, USA) for 30 min and then the oocytes were dissociated by making an incision in the abdomen. Oocytes were treated with 2 mg ml^−1^ collagenase type I (Sigma-Aldrich) for 6 hours in frog Ringer’s solution containing (mM): 88 NaCl, 1 KCl, 2.4 NaHCO_3_, 0.3 Ca (NO_3_)_2_, 0.41 CaCl_2_, 0.82 MgSO_4_, and 15 HEPES, pH 7.6 with NaOH. After removing the follicles, each oocyte was injected with 50 nl of cRNA solution and then incubated in frog Ringer’s solution with 0.1% penicillin-streptomycin (Sigma-Aldrich) at 17°C. The oocytes were used for two-electrode voltage-clamp recordings 1-5 days after the injection of cRNA.

### Two-electrode voltage-clamp recordings

Glass electrodes for two-electrode voltage-clamp recordings had a resistance of 0.2-0.5 MΩ when filled with a pipette solution containing 3 M potassium acetate and 10 mM KCl. Membrane currents of oocytes were recorded at 22-25°C in a high K^+^ solution containing (mM): 96 KCl, 3 MgCl_2_, and 5 HEPES, pH 7.3 with n-methyl-d-glucamine (NMDG). To evaluate the ion selectivity, the KCl was replaced by LiCl, NaCl, RbCl, CsCl, methylammonium-Cl, tetramethylammonium-Cl, tetraethylammonium-Cl, or NMDG-Cl, one at a time. Data were recorded by an oocyte clamp amplifier OC-725C (Warner instruments, Holliston, MA, USA), a digital converter Digidata 1440 (Molecular devices, San Jose, CA, USA), and pCLAMP 10 software (Molecular devices). Compounds were applied to the extracellular solution and perfused the oocytes in a recording chamber. Sequential applications of blockers to the extracellular solution were performed and the recording data were fitted to a Hill equation to acquire the dose-inhibition relationship as follows: y = min + (max − min) / (1 + (x/IC_50_) ^ (Hillslope)), where x refers to the concentration of blockers, y refers to the normalized current at a certain concentration, and IC_50_ is the concentration that generates half of the maximal inhibition.

### Single-channel recordings

Cell-attached recordings were carried out using *ltk-* mouse fibroblast cells as previously described (Eldstrom, Wang, Werry, Wong, & Fedida, 2015; Murray et al., 2016; Westhoff, Eldstrom, Murray, Thompson, & Fedida, 2019). Briefly, cells were transfected using Lipofectamine 2000 (Invitrogen, Carlsbad, CA, USA) as per the manufactures protocol, with DNA ratios of 1:0.8 of Kir3.2 G156S:GFP in micrograms. Transfected cells were grown in MEM with 10% FBS (GIBCO, USA), overnight in 5% CO_2_ at 37°C. All recordings were made at room temperature, ~24 hrs post transfection. The bath solution contained (mM): 135 KCl, 10 HEPES, 1 MgCl_2_, 10 Dextrose, 0.05 CaCl_2_ and was adjusted to pH 7.4 with KOH. The patch pipette solution contained (mM): 10 HEPES, 1 MgCl_2_, 10 Dextrose and either 135 KCl or 135 LiCl pH’ed to 7.4 with KOH or LiOH or a 50% mixture of each of these solutions.

Single channel recordings were made using an Axopatch 200B amplifier, Digidata 1330 and pClamp 9 software (Molecular Devices). Patch electrodes were pulled from thick-walled borosilicate glass (Sutter Instruments, Novato, CA, USA). After fire polishing and Sylgard (Dow Corning, Midland, MI, USA) application, electrode resistances were between 40 and 60 MΩ. Records were sampled at 10 kHz and low-pass filtered at 2 kHz using a −3dB, four-pole Bessel filter. For presentation and analysis, the data was additionally filtered digitally at 200 Hz. Data was analyzed with pClamp 10 (Molecular Devices) and Figures prepared using Prism 9 GraphPad software (San Diego, CA, USA).

### Data and statistical analysis

All statistical analyses were performed with SigmaPlot 14 (Hulinks, Japan). Data shown in all Figures are means ± SD from n single cells specified in the legend of each Figure. Statistical differences were evaluated using Tukey’s test following one-way ANOVA to compare the values between the absence or presence of pore blockers with the extracellular solution containing different cations in Figure 2, 3, 6, 7 and SI appendix, Figure. S1. Unpaired Student’s *t* test was used for Figures 3i, 3j and 4e.

### Reagents

Tertiapin-Q (TPN-Q) was purchased from Abcam (Cambridge, UK) or Bio-Techne Canada (Toronto, Canada). ACh, Ivermectin (IVM), and the other reagents were purchased from Sigma-Aldrich (St. Louis, USA), unless otherwise specified. TPN-Q and ACh were dissolved in distilled water, and IVM was dissolved in DMSO. These reagents were diluted to a final concentration in the extracellular solution and the solvent concentration was ≤ 0.3%.

## Acknowledgments and funding sources

We thank Ms. Chizue Naito, Ms. Tomomi Yamamoto and Fariba Ataei for technical assistance, members in Kubo Lab for discussion, and Dr. Eitan Reuveny and Dr. Izhar Karbat (Weizmann Institute of Science) for information exchange and discussion. I-S. C. thanks Dr. Tomoe Y. Nakamura (Wakayama Medical University) for support. This work was supported by JSPS KAKENHI Grant Numbers JP20K07304 (to I-S.C.) and JP20H03424 (to Y.K.), The Uehara Memorial Foundation (to I-S.C.), and grants PJT-156181 from the CIHR (to D.F.) and G-20-0028720 from the Heart and Stroke Foundation of Canada (to D.F.).

## Author Contributions

I-S.C. and Y.K. designed the study; I-S.C. performed all experiments using *Xenopus* oocytes and data analyses. J.E. performed single channel recordings and J.E. and D.F. analyzed and interpreted the data. All authors wrote the paper and confirmed the final version.

---

**Figure 1-figure supplement 1.**
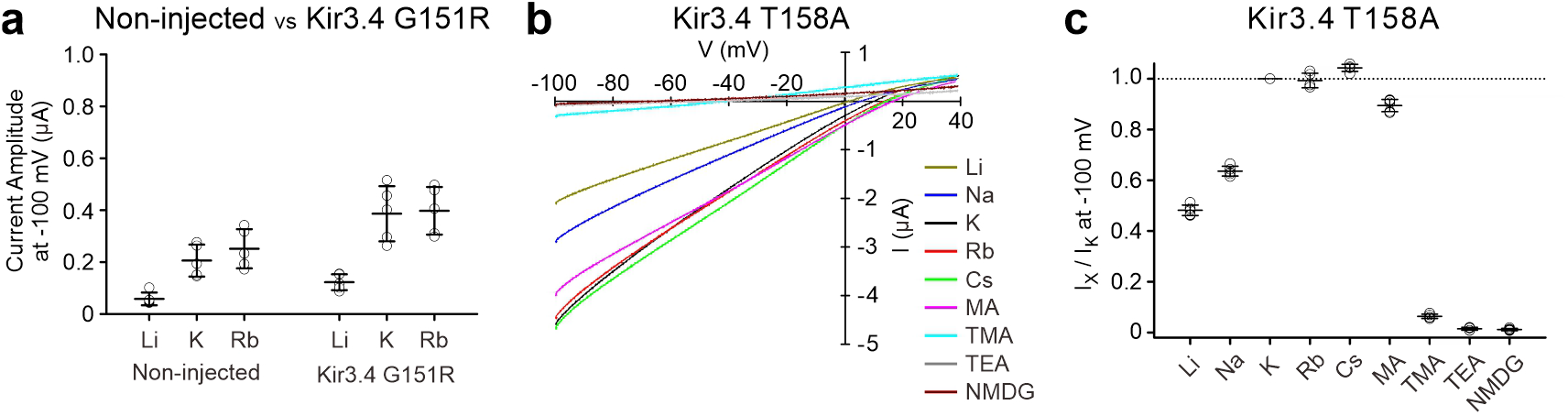
Permeation of various cations in Kir3.4 G151R and Kir3.4 T158A expressed in *Xenopus* oocytes. (**a**) The current amplitude recorded from the oocytes without injection of cRNA or injected with cRNA of Kir3.4 G151R at −100 mV in the extracellular solution containing 96 mM LiCl, KCl, or RbCl, respectively. (**b**) The IV-relationship of Kir3.4 T158A with different cations in the extracellular solution. (**c**) The current amplitude of X^+^ relative to K^+^ (I_X_/I_K_) of each cation was calculated by normalization of I_K_ as 1. Data shown are individual values with means ± SD, n = 5.

**Figure 2-figure supplement 1.**
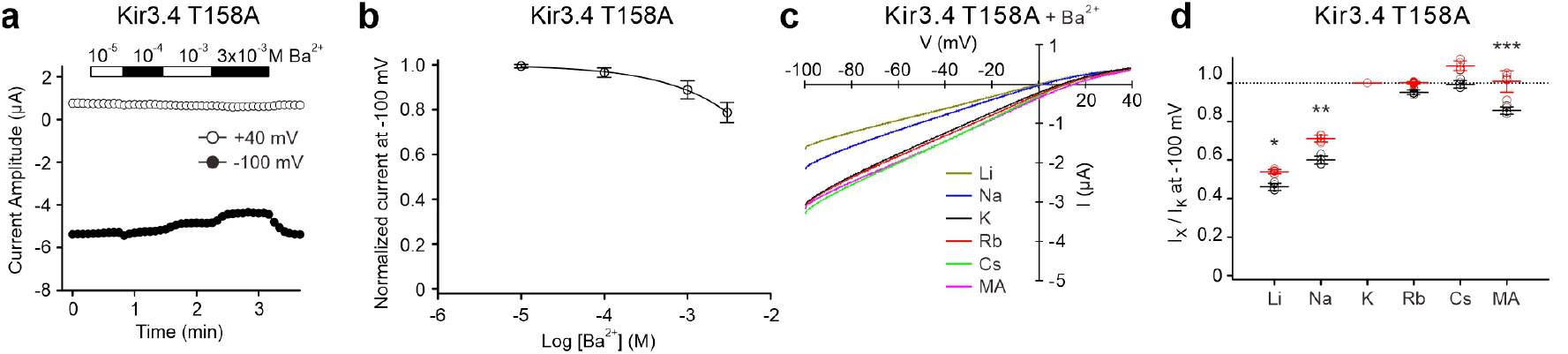
Effect of Ba^2+^ on the current of Kir3.4 T158A expressed in *Xenopus* oocytes. (**a**) Time course of current change of oocytes expressed Kir3.4 T158A at −100 mV and +40 mV before and after applications of different concentrations of Ba^2+^ in the 96 mM KCl extracellular solution. (**b**) Dose-inhibition relationship of Ba^2+^ on K^+^ current, means ± SD, n = 5. (**c**) The IV-relationship of Kir3.4 T158A in the presence of 3 mM Ba^2+^. (**d**) The I_X_/I_K_ at −100 mV before (black symbols) and after the application of 3 mM Ba^2+^ (red symbols), means ± SD, n = 5. * *P* < 0.05, ** *P* < 0.01, *** *P* < 0.001 depict the statistical significance of the difference between the absence and the presence of Ba^2+^.

**Figure 5-figure supplement 1.**
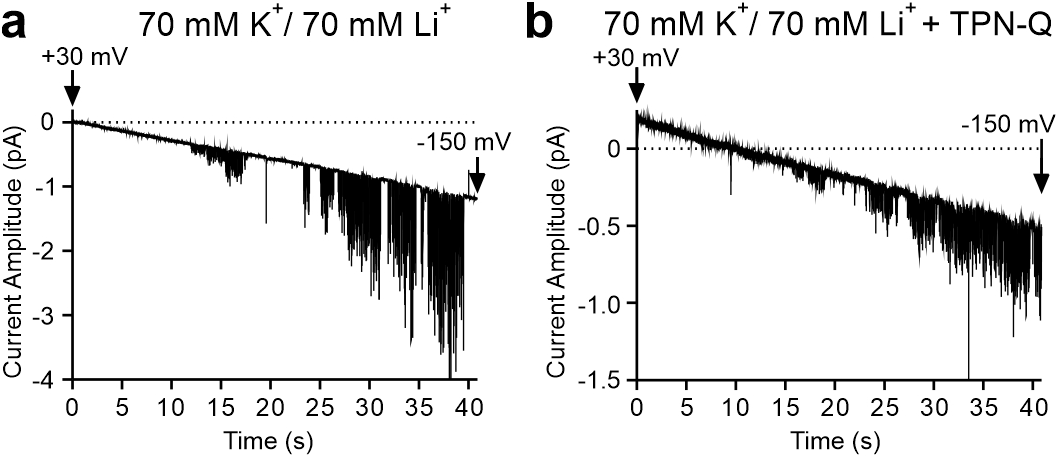
The voltage-conductance relationship of Kir3.2 G156S expressed in mouse fibroblasts. Single-channel recordings of Kir3.2 G156S were recorded with pipette solution containing 70 mM K^+^ and 70 mM Li^+^ in the absence (**a**) or presence of 400 nM TPN-Q (**b**) with a ramp pulse from +30 mV to −150 mV for 40 s.

**Figure 7-figure supplement 1.**
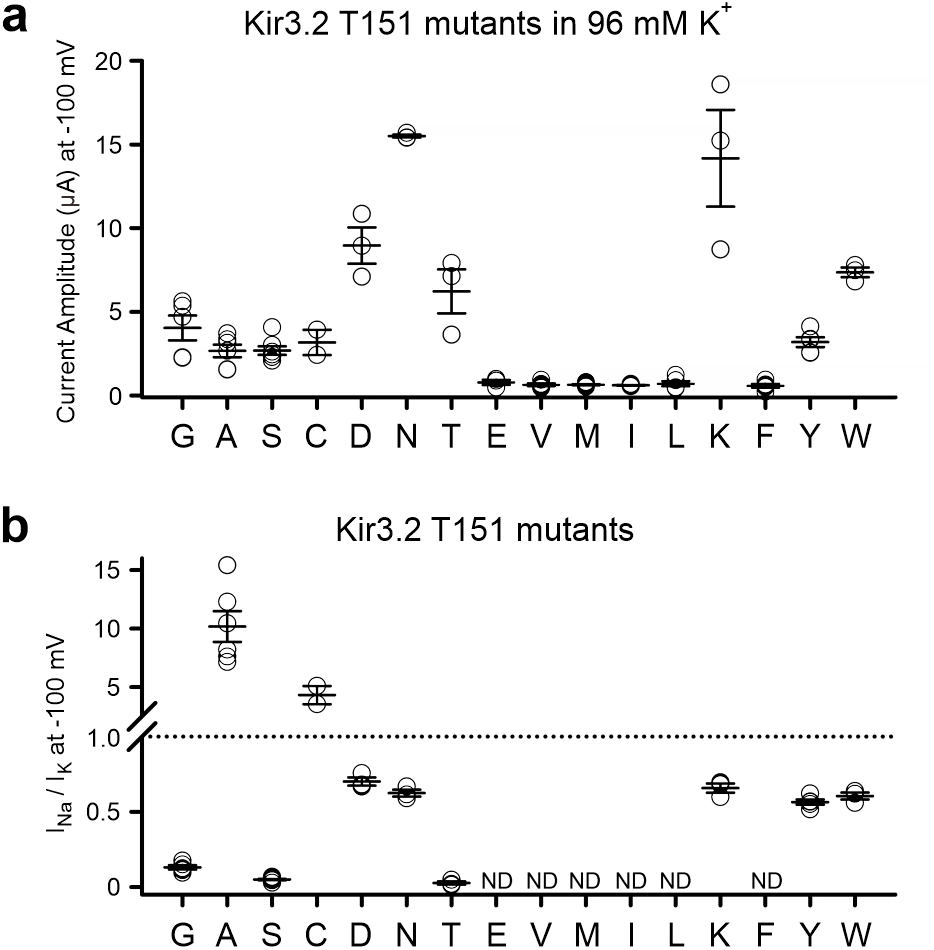
The current amplitude and the ion selectivity of various Kir3.2 mutants at T151 expressed in *Xenopus* oocytes. (**a**) The basal current amplitude of Kir3.2 T151 mutants at −100 mV in the 96 mM KCl extracellular solution. (**b**) The I_Na_/I_K_ of Kir3.2 T151 mutants at −100 mV. All data shown are means ± SD, n = 5 for T151G, V, L, F, Y; n = 6 for T151A, I; n = 7 for T151S; n = 8 for T151M; n = 2 for T151C; n = 3 for other mutants. The I_Na_/I_K_ of T151E, V, M, I, L, F mutants is not detectable (indicated as ND) because of their very small current amplitudes as shown in (**a**).

**Figure 7-figure supplement 2.**
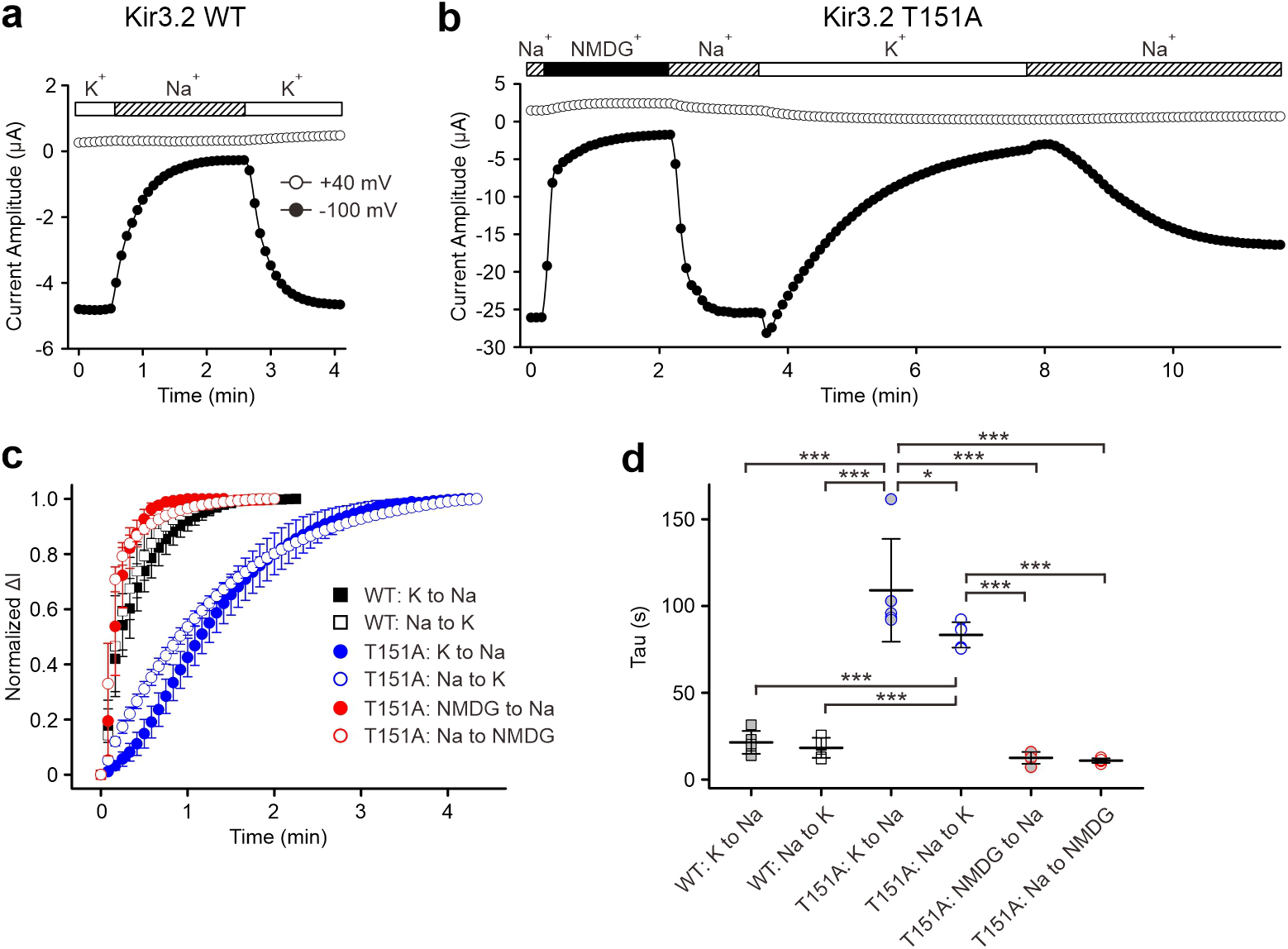
The kinetics of current amplitude change of Kir3.2 T151A upon exchange of extracellular cations. (**a, b**) Representative time course of current amplitude change of Kir3.2 WT (**a**) and Kir3.2 T151A mutant (**b**) at −100 mV and +40 mV during the exchange of extracellular solution with 96 mM K^+^ or Na^+^ or NMDG^+^ as shown above. (**c**) The kinetics of current amplitude change upon the exchange of extracellular cations calculated from the current amplitude at −100 mV. The maximal difference of current amplitude before and after replacement of extracellular solution was normalized to 1. Data shown are means ± SD, n = 5 for each. (**d**) Tau of current amplitude change, means ± SD, n = 5 for each. * *P* < 0.05, *** *P* < 0.001 depict the statistical significance.

## References

Alagem, N., Dvir, M., & Reuveny, E. (2001). Mechanism of Ba(2+) block of a mouse inwardly rectifying K+ channel: differential contribution by two discrete residues. J Physiol, 534(Pt. 2), 381–393. doi:10.1111/j.1469-7793.2001.00381.x

Bichet, D., Grabe, M., Jan, Y. N., & Jan, L. Y. (2006). Electrostatic interactions in the channel cavity as an important determinant of potassium channel selectivity. Proc Natl Acad Sci U S A, 103(39), 14355–14360. doi:10.1073/pnas.0606660103

Bichet, D., Lin, Y. F., Ibarra, C. A., Huang, C. S., Yi, B. A., Jan, Y. N., & Jan, L. Y. (2004). Evolving potassium channels by means of yeast selection reveals structural elements important for selectivity. Proc Natl Acad Sci U S A, 101(13), 4441–4446. doi:10.1073/pnas.0401195101

Boulkroun, S., Beuschlein, F., Rossi, G. P., Golib-Dzib, J. F., Fischer, E., Amar, L., … Zennaro, M. C. (2012). Prevalence, clinical, and molecular correlates of KCNJ5 mutations in primary aldosteronism. Hypertension, 59(3), 592–598. doi:10.1161/HYPERTENSIONAHA.111.186478

Chen, I. S., & Kubo, Y. (2018). Ivermectin and its target molecules: shared and unique modulation mechanisms of ion channels and receptors by ivermectin. J Physiol, 596(10), 1833–1845. doi:10.1113/JP275236

Chen, I. S., Liu, C., Tateyama, M., Karbat, I., Uesugi, M., Reuveny, E., & Kubo, Y. (2019). Non-sedating antihistamines block G-protein-gated inwardly rectifying K(+) channels. Br J Pharmacol, 176(17), 3161–3179. doi:10.1111/bph.14717

Chen, I. S., Tateyama, M., Fukata, Y., Uesugi, M., & Kubo, Y. (2017). Ivermectin activates GIRK channels in a PIP2 -dependent, Gbetagamma -independent manner and an amino acid residue at the slide helix governs the activation. J Physiol, 595(17), 5895–5912. doi:10.1113/JP274871

Choi, M., Scholl, U. I., Yue, P., Bjorklund, P., Zhao, B., Nelson-Williams, C., … Lifton, R. P. (2011). K+ channel mutations in adrenal aldosterone-producing adenomas and hereditary hypertension. Science, 331(6018), 768–772. doi:10.1126/science.1198785

Doupnik, C. A., Parra, K. C., & Guida, W. C. (2015). A computational design approach for virtual screening of peptide interactions across K(+) channel families. Comput Struct Biotechnol J, 13, 85–94. doi:10.1016/j.csbj.2014.11.004

Doyle, D. A., Morais Cabral, J., Pfuetzner, R. A., Kuo, A., Gulbis, J. M., Cohen, S. L., … MacKinnon, R. (1998). The structure of the potassium channel: molecular basis of K+ conduction and selectivity. Science, 280(5360), 69–77. doi:10.1126/science.280.5360.69

Eisenman, G. (1962). Cation selective glass electrodes and their mode of operation. Biophys J, 2(2 Pt 2), 259–323. doi:10.1016/s0006-3495(62)86959-8

Eldstrom, J., Wang, Z., Werry, D., Wong, N., & Fedida, D. (2015). Microscopic mechanisms for long QT syndrome type 1 revealed by single-channel analysis of I(Ks) with S3 domain mutations in KCNQ1. Heart Rhythm, 12(2), 386–394. doi:10.1016/j.hrthm.2014.10.029

Guo, R., Zeng, W., Cui, H., Chen, L., & Ye, S. (2014). Ionic interactions of Ba2+ blockades in the MthK K+ channel. J Gen Physiol, 144(2), 193–200. doi:10.1085/jgp.201411192

Heginbotham, L., Lu, Z., Abramson, T., & MacKinnon, R. (1994). Mutations in the K+ channel signature sequence. Biophys J, 66(4), 1061–1067. doi:10.1016/S0006-3495(94)80887-2

Hibino, H., Inanobe, A., Furutani, K., Murakami, S., Findlay, I., & Kurachi, Y. (2010). Inwardly rectifying potassium channels: their structure, function, and physiological roles. Physiol Rev, 90(1), 291–366. doi:10.1152/physrev.00021.2009

Hille, B. (2001). Sinauer Associates Inc.

Horvath, G. A., Zhao, Y., Tarailo-Graovac, M., Boelman, C., Gill, H., Shyr, C., … van Karnebeek, C. D. M. (2018). Gain-of-function KCNJ6 Mutation in a Severe Hyperkinetic Movement Disorder Phenotype. Neuroscience, 384, 152–164. doi:10.1016/j.neuroscience.2018.05.031

Hsieh, C. P., Kuo, C. C., & Huang, C. W. (2015). Driving force-dependent block by internal Ba(2+) on the Kir2.1 channel: Mechanistic insight into inward rectification. Biophys Chem, 202, 40–57. doi:10.1016/j.bpc.2015.04.003

Kofuji, P., Hofer, M., Millen, K. J., Millonig, J. H., Davidson, N., Lester, H. A., & Hatten, M. E. (1996). Functional analysis of the weaver mutant GIRK2 K+ channel and rescue of weaver granule cells. Neuron, 16(5), 941–952. doi:10.1016/s0896-6273(00)80117-8

Krapivinsky, G., Gordon, E. A., Wickman, K., Velimirovic, B., Krapivinsky, L., & Clapham, D. E. (1995). The G-protein-gated atrial K+ channel IKACh is a heteromultimer of two inwardly rectifying K(+)-channel proteins. Nature, 374(6518), 135–141. doi:10.1038/374135a0

Kubo, Y., Reuveny, E., Slesinger, P. A., Jan, Y. N., & Jan, L. Y. (1993). Primary structure and functional expression of a rat G-protein-coupled muscarinic potassium channel. Nature, 364(6440), 802–806. doi:10.1038/364802a0

Lesage, F., Duprat, F., Fink, M., Guillemare, E., Coppola, T., Lazdunski, M., & Hugnot, J. P. (1994). Cloning provides evidence for a family of inward rectifier and G-protein coupled K+ channels in the brain. FEBS Lett, 353(1), 37–42. doi:10.1016/0014-5793(94)01007-2

Long, S. B., Campbell, E. B., & Mackinnon, R. (2005). Crystal structure of a mammalian voltage-dependent Shaker family K+ channel. Science, 309(5736), 897–903. doi:10.1126/science.1116269

Maiolino, G., Ceolotto, G., Battistel, M., Barbiero, G., Cesari, M., Amar, L., … Rossi, G. P. (2018). Macrolides for KCNJ5-mutated aldosterone-producing adenoma (MAPA): design of a study for personalized diagnosis of primary aldosteronism. Blood Press, 27(4), 200–205. doi:10.1080/08037051.2018.1436961

Masotti, A., Uva, P., Davis-Keppen, L., Basel-Vanagaite, L., Cohen, L., Pisaneschi, E., … Dallapiccola, B. (2015). Keppen-Lubinsky syndrome is caused by mutations in the inwardly rectifying K+ channel encoded by KCNJ6. Am J Hum Genet, 96(2), 295–300. doi:10.1016/j.ajhg.2014.12.011

Matamoros, M., & Nichols, C. G. (2021). Pore-forming transmembrane domains control ion selectivity and selectivity filter conformation in the KirBac1.1 potassium channel. J Gen Physiol, 153(5). doi:10.1085/jgp.202012683

Miller, A. N., & Long, S. B. (2012). Crystal structure of the human two-pore domain potassium channel K2P1. Science, 335(6067), 432–436. doi:10.1126/science.1213274

Murray, C. I., Westhoff, M., Eldstrom, J., Thompson, E., Emes, R., & Fedida, D. (2016). Unnatural amino acid photo-crosslinking of the IKs channel complex demonstrates a KCNE1:KCNQ1 stoichiometry of up to 4:4. Elife, 5. doi:10.7554/eLife.11815

Navarro, B., Kennedy, M. E., Velimirovic, B., Bhat, D., Peterson, A. S., & Clapham, D. E. (1996). Nonselective and G betagamma-insensitive weaver K+ channels. Science, 272(5270), 1950–1953. doi:10.1126/science.272.5270.1950

Patel, D., Kuyucak, S., & Doupnik, C. A. (2020). Structural Determinants Mediating Tertiapin Block of Neuronal Kir3.2 Channels. Biochemistry, 59(7), 836–850. doi:10.1021/acs.biochem.9b01098

Patil, N., Cox, D. R., Bhat, D., Faham, M., Myers, R. M., & Peterson, A. S. (1995). A potassium channel mutation in weaver mice implicates membrane excitability in granule cell differentiation. Nat Genet, 11(2), 126–129. doi:10.1038/ng1095-126

Prutting, S., & Grissmer, S. (2011). A novel current pathway parallel to the central pore in a mutant voltage-gated potassium channel. J Biol Chem, 286(22), 20031–20042. doi:10.1074/jbc.M110.185405

Roy, R. N., Hendriks, K., Kopec, W., Abdolvand, S., Weiss, K. L., de Groot, B. L., … Coates, L. (2021). Structural plasticity of the selectivity filter in a nonselective ion channel. IUCrJ, 8(Pt 3), 421–430. doi:10.1107/S205225252100213X

Scholl, U. I., Abriola, L., Zhang, C., Reimer, E. N., Plummer, M., Kazmierczak, B. I., … Lifton, R. P. (2017). Macrolides selectively inhibit mutant KCNJ5 potassium channels that cause aldosterone-producing adenoma. J Clin Invest, 127(7), 2739–2750. doi:10.1172/JCI91733

Scholl, U. I., Nelson-Williams, C., Yue, P., Grekin, R., Wyatt, R. J., Dillon, M. J., … Lifton, R. P. (2012). Hypertension with or without adrenal hyperplasia due to different inherited mutations in the potassium channel KCNJ5. Proc Natl Acad Sci U S A, 109(7), 2533–2538. doi:10.1073/pnas.1121407109

Shi, C., He, Y., Hendriks, K., de Groot, B. L., Cai, X., Tian, C., … Sun, H. (2018). A single NaK channel conformation is not enough for non-selective ion conduction. Nat Commun, 9(1), 717. doi:10.1038/s41467-018-03179-y

Slesinger, P. A., Patil, N., Liao, Y. J., Jan, Y. N., Jan, L. Y., & Cox, D. R. (1996). Functional effects of the mouse weaver mutation on G protein-gated inwardly rectifying K+ channels. Neuron, 16(2), 321–331. doi:10.1016/s0896-6273(00)80050-1

Tyutyaev, P., & Grissmer, S. (2017). Observation of sigma-pore currents in mutant hKv1.2_V370C potassium channels. PLoS One, 12(4), e0176078. doi:10.1371/journal.pone.0176078

Tyutyaev, P., & Grissmer, S. (2018). Characterization of the sigma-Pore in Mutant hKv1.3 Potassium Channels. Cell Physiol Biochem, 46(3), 1112–1121. doi:10.1159/000488840

Velarde-Miranda, C., Gomez-Sanchez, E. P., & Gomez-Sanchez, C. E. (2013). Regulation of aldosterone biosynthesis by the Kir3.4 (KCNJ5) potassium channel. Clin Exp Pharmacol Physiol, 40(12), 895–901. doi:10.1111/1440-1681.12151

Wang, S., Lee, S. J., Maksaev, G., Fang, X., Zuo, C., & Nichols, C. G. (2019). Potassium channel selectivity filter dynamics revealed by single-molecule FRET. Nat Chem Biol, 15(4), 377–383. doi:10.1038/s41589-019-0240-7

Wang, W., & MacKinnon, R. (2017). Cryo-EM Structure of the Open Human Ether-a-go-go-Related K(+) Channel hERG. Cell, 169(3), 422–430 e410. doi:10.1016/j.cell.2017.03.048

Westhoff, M., Eldstrom, J., Murray, C. I., Thompson, E., & Fedida, D. (2019). I Ks ion-channel pore conductance can result from individual voltage sensor movements. Proc Natl Acad Sci U S A, 116(16), 7879–7888. doi:10.1073/pnas.1811623116

Whorton, M. R., & MacKinnon, R. (2011). Crystal structure of the mammalian GIRK2 K+ channel and gating regulation by G proteins, PIP2, and sodium. Cell, 147(1), 199–208. doi:10.1016/j.cell.2011.07.046

Yang, J., Yu, M., Jan, Y. N., & Jan, L. Y. (1997). Stabilization of ion selectivity filter by pore loop ion pairs in an inwardly rectifying potassium channel. Proc Natl Acad Sci U S A, 94(4), 1568–1572. doi:10.1073/pnas.94.4.1568

Yi, B. A., Lin, Y. F., Jan, Y. N., & Jan, L. Y. (2001). Yeast screen for constitutively active mutant G protein-activated potassium channels. Neuron, 29(3), 657–667. doi:10.1016/s0896-6273(01)00241-0

Zagotta, W. N. (2006). Membrane biology: permutations of permeability. Nature, 440(7083), 427–429. doi:10.1038/440427a

Zangerl-Plessl, E. M., Qile, M., Bloothooft, M., Stary-Weinzinger, A., & van der Heyden, M. A. G. (2019). Disease Associated Mutations in KIR Proteins Linked to Aberrant Inward Rectifier Channel Trafficking. Biomolecules, 9(11). doi:10.3390/biom9110650

Zhou, Y., Morais-Cabral, J. H., Kaufman, A., & MacKinnon, R. (2001). Chemistry of ion coordination and hydration revealed by a K+ channel-Fab complex at 2.0 A resolution. Nature, 414(6859), 43–48. doi:10.1038/35102009

