## Supplementary Information for "A novel ion conducting route besides the central pore in an inherited mutant of G-protein-gated inwardly rectifying K^+^ channel"

### Supplementary Table 1

| Mutant | Forward primer (5’→3’) | Reversed primer (3’→5’) |
|---|---|---|
| Kir3.2 | | |
| S148F | gcttttttattcttcatagagacagaaacc | ggtttctgtctctatgaagaataaaaaagc |
| T151G | ttctccatagagggagaaaccacca | tggtggtttctccctctatggagaa |
| T151A | ttctccatagaggcagaaaccacc | ggtggtttctgcctctatggagaa |
| T151S | ttctccatagagtcagaaaccacc | ggtggtttctgactctatggagaa |
| T151C | ttctccatagagtgcgaaaccaccatc | gatggtggtttcgcactctatggagaa |
| T151D | ttctccatagaggacgaaaccaccatc | gatggtggtttcgtcctctatggagaa |
| T151N | ttctccatagagaacgaaaccaccatc | gatggtggtttcgttctctatggagaa |
| T151E | tctccatagaggaagaaaccacca | tggtggtttcttcctctatggaga |
| T151V | ttctccatagaggtagaaaccacca | tggtggtttctacctctatggagaa |
| T151M | ttctccatagagatggaaaccaccatc | gatggtggtttccatctctatggagaa |
| T151I | ttctccatagagatagaaaccacca | tggtggtttctatctctatggagaa |
| T151L | ttctccatagagctagaaaccacca | tggtggtttctagctctatggagaa |
| T151K | ttctccatagagaaagaaaccacc | ggtggtttctttctctatggagaa |
| T151F | ttctccatagagttcgaaaccaccatc | gatggtggtttcgaactctatggagaa |
| T151Y | ttctccatagagtacgaaaccaccatc | gatggtggtttcgtactctatggagaa |
| T151W | ttctccatagagtgggaaaccaccatc | gatggtggtttcccactctatggagaa |
| T154del | atcggttatggctaccgggtcatca | ggtttctgtctctatggagaataa |
| G156S | gaaaccaccatctcttatggctac | gtagccataagagatggtggtttc |
| G156R | gaaaccaccatccgttatggctac | gtagccataacggatggtggtttc |
| T163A | taccgggtcatcgcggacaagtg | cacttgtccgcgatgacccggta |
| L173R | gggattattctccgcttaatccag | ctggattaagcggagaataatccc |
| Kir3.4 | | |
| G151R | gaaacaaccattcggtatggcttc | gaagccataccgaatggttgtttc |
| T158A | tcagagtcattgcagagaagtgtc | gacacttctctgcaatgactctga |

**Supplementary Table 1. Primer sequences**

The primers used for constructions of Kir3.2 and Kir3.4 mutants by site-directed mutagenesis using PfuUltra II Fusion HS DNA Polymerase kit (Agilent, USA).
